# Ser129 Phosphorylation Paradox: Non-phosphorylated α-Synuclein Drives Parkinson’s Disease-like Pathogenesis *in Vivo*

**DOI:** 10.64898/2025.12.21.695762

**Authors:** Zhengming Tian, Zirui Xu, Mengyuan Guo, Haoyi Fang, Huiyi Yang, Qianqian Shao, Dongjun Wang, Feiyang Jin, Yakun Gu, Guiyou Liu, Rehana K Leak, Jun Chen, Jia Liu

**Affiliations:** Beijing Institute of Brain Disorders, Laboratory of Brain Disorders, Hypoxia Conditioning Translational Laboratory of Clinical Medicine, Ministry of Science and Technology, Chinese Institutes for Medical Research, Collaborative Innovation Center for Brain Disorders, Capital Medical University, Beijing, China; Pittsburgh Institute of Brain Disorders and Recovery and Department of Neurology, University of Pittsburgh School of Medicine, Pittsburgh, PA, USA; Graduate School of Pharmaceutical Sciences, Duquesne University, 418C Mellon Hall, 913 Bluff Street, Pittsburgh, PA, 15219, USA

## Abstract

Accumulation of phosphorylated α-synuclein at serine 129 is a hallmark of Parkinson’s disease (PD), but its pathogenic role remains controversial. We conducted longitudinal studies using non-phosphorylatable S129A and phosphomimetic S129D α-synuclein global knock-in mice. Unexpectedly, the S129A mice—rather than S129D mice—developed reproducible olfactory, motor, and cognitive deficits, dopaminergic dysfunction and neurodegeneration, α-synuclein insolubility, and Lewy-like inclusions by 3 to 9 months, with progressive worsening up to 12 months of age. Spatial transcriptomics and electron microscopy revealed mitochondrial, synaptic, and protein ubiquitination abnormalities in the substantia nigra of S129A α-synuclein mice. Viral vector-mediated nigral overexpression of S129D-α-synuclein or systemic administration of an anti-α-synuclein-aggregation drug partially alleviated the early neurological phenotypes of S129A mice. These results implicate non-phosphorylatable α-synuclein as a direct pathogenic driver for PD.

## Main Text

Parkinson’s disease (PD), the second most common neurodegenerative disorder, is manifested through progressive motor deficits and non-motor symptoms spanning prodromal to advanced stages(*1, 2*). At the histopathological level, PD is defined by nigral dopaminergic neuron loss and Lewy bodies (LBs)—intracellular inclusions dominated by α-synuclein (α-syn) phosphorylated at serine 129 (pS129)(*3, 4*). pS129 accounts for >90% of the aggregated α-syn in LBs versus a the vastly smaller fraction in normal brains, a striking association that has established α-syn hyperphosphorylation as a major pathologic hallmark(*5, 6*). However, the mechanistic role of phosphorylation at this residue remains unresolved: Does this modification drive neurodegeneration, protect neurons, or simply mark disease progression?

Emerging evidence complicates this paradigm. Physiological increases in α-syn pS129 during cellular stress suggest potential functional roles (*7–9*), but the *in vivo* properties of phosphorylated versus non-phosphorylated α-syn remain unresolved. To address this fundamental gap, we developed S129A (non-phosphorylatable) and S129D (phosphomimetic) α-syn knock-in (KI) mice. Contrary to our initial expectations, S129D mice demonstrated little to no PD-related pathology. Surprisingly, the S129A mice better recapitulated human PD progression—including temporally stratified motor/non-motor (olfactory, intestinal, and cognitive) deficits and nigrostriatal dopaminergic degeneration—whereas the S129D phenotype only showed intestinal dysfunction. Spatial transcriptomics further linked non-phosphorylated α-syn to pathological processes related to PD. These findings overturned our conventional assumptions about the pathogenic role of pS129 and implicated non-phosphorylated rather than phosphorylated α-syn as the potential driver of PD-like pathologies, providing us with a uniquely relevant genetic model for translational research.

### Progressive motor deficits and nigrostriatal dopaminergic degeneration in S129A KI mice

To dissect the role of pS129 α-syn in PD pathogenesis, we generated S129A (non-phosphorylatable) and S129D (phosphomimetic) *Snca* KI mice via CRISPR–Cas9 (Fig. S1A), which preserves endogenous levels of α-syn expression and avoids overexpression artifacts. Genotyping and sequencing confirmed the S129 mutations (Fig. S1B). Western blotting confirmed equivalent total α-syn levels across genotypes. Neither KI line had detectable pS129 α-syn because S129A mimics the non-phosphorylated charge state while S129D mimics the phosphorylated charge state(*10*). The two KI lines had comparable β-syn levels, suggesting lack of compensatory induction of this member of the synuclein family (Fig. S1C–F). Additionally, S129A/S129D KI mice exhibited normal body morphologies, weaning survival rates, food intake, body weights, and organ histopathology (Fig. S1G–L)—excluding overt, non-specific artifacts.

We assessed PD-relevant phenotypes in WT, S129A KI, and S129D KI mice across 1–12 months (M), focusing on motor function, dopaminergic (DA) neuron histology, and DA metabolism (Fig. 1A). First, we evaluated motor function using the Rotarod and pole tests: starting at 6 M, S129A KI mice showed significantly longer descent latencies in the pole test (Fig. 1B, C) and progressively shorter fall latencies in the Rotarod (Fig. 1D, E) with a trend towards progressive worsening. Paired analyses further revealed marked motor decline in S129A KI mice between 3 and 6 M (Fig. 1F, G). In contrast, S129D KI mice performed indistinguishably from WT mice. In integrating thepole test and Rotarod data with a principal component analysis, the WT and S129D KI mice clustered relatively tightly across 3–12 M, indicating stable motor performances; however, S129A KI mice progressively diverged from this cluster with age, supporting a PD-like decay in motor function (Fig. 1H).

**Fig. 1.**
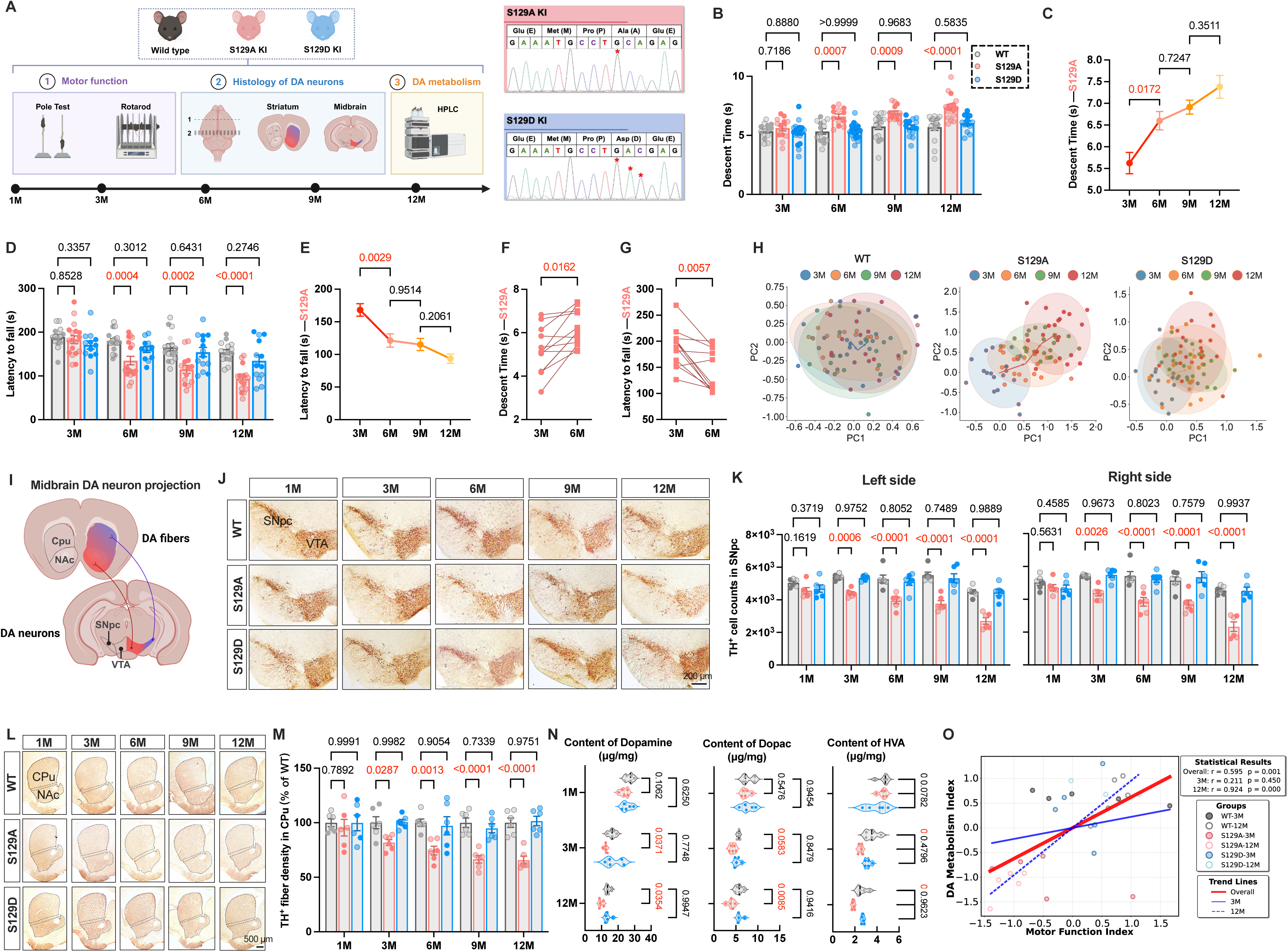
S129A, but not S129D, knock-in mice develop progressive PD-like motor deficits and nigrostriatal dopaminergic degeneration. (A) Schematic of the experimental design: PD-relevant phenotypes, including motor function, DA neuron histology, and DA metabolism, were assessed in WT, S129A KI, and S129D KI mice across 1–12 months (M). Right panels show sequencing validation of S129A/S129D site mutations in the *Snca* gene. (B) Pole Test descent latencies across genotypes and ages. n = 12 mice per group, including 6 males (dark points) and 6 females (light points). (C) Age-related longitudinal progression of Pole Test descent latencies in S129A KI mice. (D) Rotarod fall latencies across genotypes and ages. n = 12 mice per group, including 6 males and 6 females. (E) Age-related longitudinal progression of Rotarod fall latencies in S129A KI mice. (F–G) Paired analyses of individual mice, showing motor decline (Pole Test: F; Rotarod: G) between 3 and 6 months. (H) Principal component analysis integrating Pole Test and Rotarod data across genotypes and ages. Each point represents an individual mouse; ellipses indicate 95% confidence intervals. (I) Schematic of midbrain dopaminergic neuron projections: SNpc→CPu; VTA →NAc. (J–K) TH (dopaminergic neuron marker) immunostaining of SNpc/VTA in midbrain across genotypes and ages (J, scale bar: 200 μm); quantification of TH-positive neuron counts in left/right SNpc via stereology (K). n = 6 mice per group, including 3 males and 3 females. (L–M) TH immunostaining of CPu/NAc in striatum across genotypes and ages (L); quantification of TH-positive fiber density in CPu (normalized to WT) (M). n = 6 mice per group, including 3 males and 3 females. (N) Striatal DA, Dopac, and HVA levels quantified via high-performance liquid chromatography. n = 5 male mice per group. (O) Correlation between composite Motor Function Index and DA Metabolism Index: overall (r = 0.595, p = 0.001), 3 M (r = 0.211, p = 0.450), and 12 M (r = 0.924, p < 0.001). Each point represents an individual mouse; trend lines denote linear associations. All data are presented as mean ± SEM. Motor function analyses (B, D), histology analyses (K, M), and DA metabolism analysis (N) used two-way ANOVA with Tukey’s post hoc test. Longitudinal trend analyses of S129A KI mice (C, E) used one-way ANOVA. Paired analyses (F–G) used two-tailed paired t-test. Correlation analysis (O) used two-tailed Pearson correlation. DA, dopaminergic; DOPAC, 3,4-dihydroxyphenylacetic acid; HVA, homovanillic acid; HPLC, high-performance liquid chromatography; KI, knock-in; PCA, principal component analysis; CPu, caudate-putamen; NAc, nucleus accumbens; SNpc, substantia nigra pars compacta; TH, tyrosine hydroxylase; VTA, ventral tegmental area; WT, wild-type.

To link motor deficits to nigrostriatal pathology, we used the canonical catecholaminergic cell marker tyrosine hydroxylase (TH) alongside TH^+^NeuN^+^ co-immunostaining. A positive correlation between TH^+^NeuN^+^ and TH^+^ cell counts in the SNpc was noted (Fig. S2), suggesting that TH staining was appropriate for quantifying DA neuron loss. Then, we quantified TH^+^ neurons in the midbrain and their projections in the striatum (Fig. 1I). Relative to WT mice, S129A KI mice displayed age-dependent loss of TH^+^ neurons in the substantia nigra pars compacta (SNpc) starting at 3 M (Fig. 1J–K, S3B, F). For SNpc projections to the caudoputamen (CPu), S129A KI mice showed progressive loss of TH^+^ fibers in the dorsal striata (Fig. 1L–M, S3A, G). In contrast S129D KI mice matched WT mice in all TH staining metrics. Consistent with PD-characteristic, SNpc-selective degeneration, TH^+^ neurons in the ventral tegmental area (VTA) and the mesolimbic projections to the nucleus accumbens (NAc) remained unaltered across genotypes until 12 M, when S129A KI mice showed moderate loss of TH^+^ neurons and fibers (Fig. S3A–E). This pattern mirrored the milder VTA neuron loss in late-stage PD patients(*11, 12*).

We further assessed striatal DA content via HPLC. S129A KI mice exhibited age-dependent reductions in DA and its metabolites—3,4-dihydroxyphenylacetic acid (DOPAC) and homovanillic acid (HVA)—a hallmark dopaminergic alteration in PD (Fig. 1N). DA turnover-related metrics showed no consistent differences across genotypes (Fig. S3H–J). S129D KI mice maintained WT-equivalent DA and metabolite levels. We further integrated DA metabolic data with motor phenotypes, and a strong positive correlation between these variables confirmed coordinated worsening of motor and dopaminergic deficits in S129A KI mice (Fig. 1O).

Collectively, S129A KI mice recapitulated core hallmarks of PD: progressive motor dysfunction, nigrostriatal followed by mesolimbic DA neuron degeneration, and DA metabolic impairment. In contrast, S129D KI mice lacked such phenotypes, challenging the canonical model that pS129 α-syn drives PD pathogenesis, and implicating non-phosphorylated S129 α-syn as far more instrumental.

### Age-dependent aggregation of non-phosphorylated S129 **α**-syn in the substantia nigra

Given that S129A (but not S129D) KI mice recapitulate PD-like DA neurodegeneration, we next investigated whether S129 dephosphorylation drives pathological α-syn aggregation in the ventral midbrain. We analyzed α-syn solubility via Western blot, separating RIPA-soluble and insoluble fractions. At 3 M, RIPA-soluble α-syn levels were comparable among WT, S129A, and S129D mice, but by 12 M, S129A mice had lower levels of soluble α-syn, whereas S129D mice maintained levels similar to WT. Conversely, in the RIPA-insoluble fraction, S129A mice matched WT at 3 M but showed marked elevation by 12 M, whereas S129D mice mirrored WT at both ages (Fig. 2A–B). These results indicate age-dependent α-syn homeostasis disruption in S129A KI mice, with a shift from the soluble to insoluble fraction in the ventral midbrain.

**Fig. 2.**
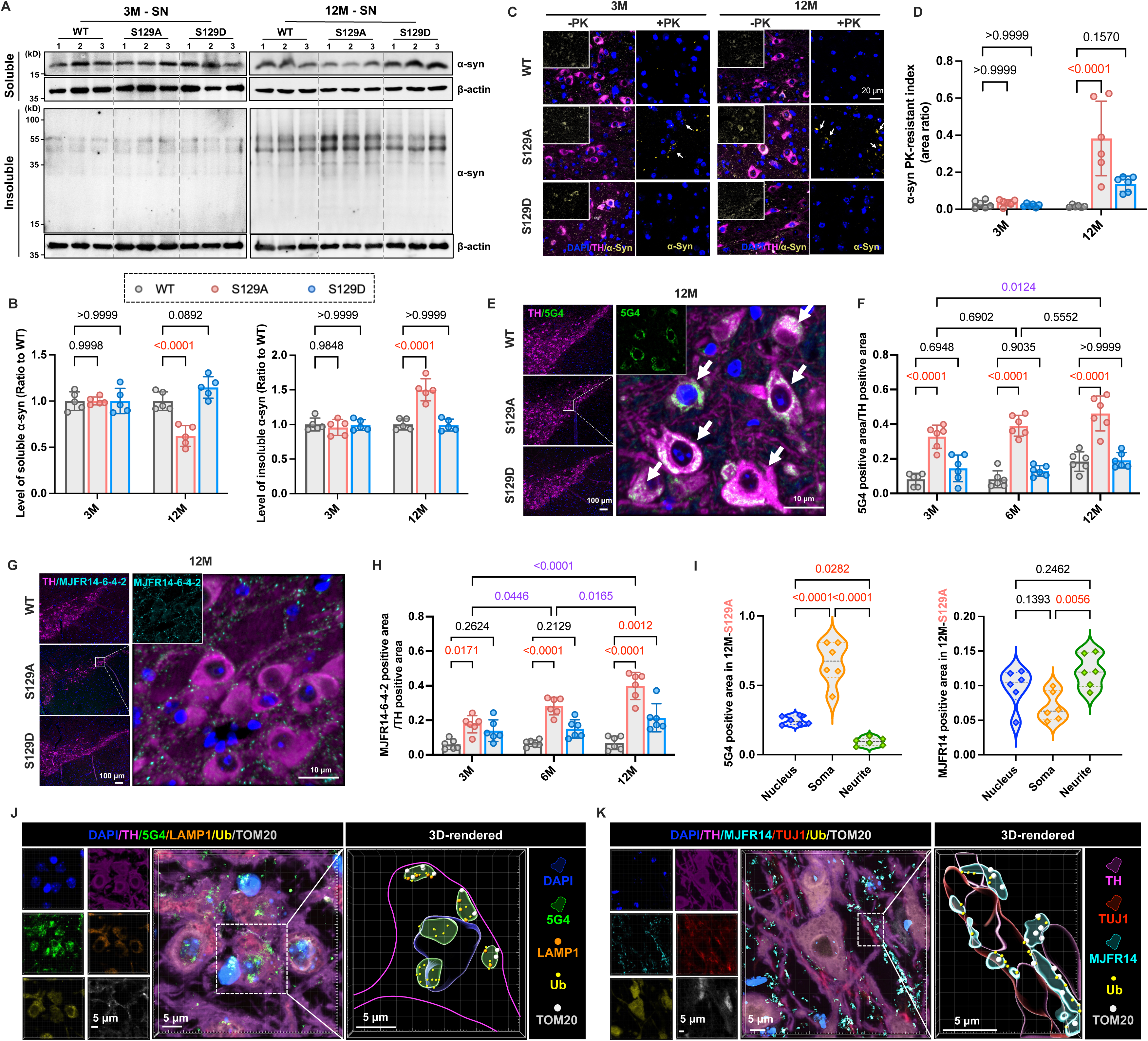
Age-dependent aggregation of non-phosphorylated S129 α-syn in the substantia nigra of S129A knock-in mice. (A–B) Western blot analysis of α-syn in RIPA-soluble and urea-solubilized RIPA-insoluble fractions of SN lysates from WT, S129A KI, and S129D KI mice at 3 and 12 M (A; β-actin serves as a loading control), and quantification of soluble (left) and insoluble (right) α-syn levels (normalized to WT) (B, n=5). (C–D) α-syn immunofluorescence staining (α-syn, yellow; TH, pink; DAPI, blue) in SN sections of WT, S129A KI, and S129D KI mice at 3 and 12M with or without proteinase K(PK) treatment, white arrows indicate PK-resistant α-syn signals (C, scale bars: 20 μm), and quantification of the α-syn PK-resistant index (area ratio in TH^+^ cell) (D, n=6). (E–F) TH (pink)/5G4 (green, aggregated α-syn) immunofluorescence co-staining in SN sections of WT, S129A KI, and S129D KI mice, white arrows indicate 5G4^+^ aggregates in TH^+^ neurons (E; 12M shown; scale bars: 100 and 10 μm), and quantification of 5G4^+^ positive area (normalized to TH^+^ area) across 3, 6, and 12M (F, n=6). (G–H) TH (pink)/MJFR14-6-4-2 (green, pathological α-syn conformation) immunofluorescence co-staining in SN sections of WT, S129A KI, and S129D KI mice (G; 12M shown; scale bars: 100 and 10 μm) and quantification of MJFR14-6-4-2^+^ positive area (normalized to TH^+^ area) across 3, 6, and 12M (H, n=6). (I) Subcellular localization analysis of 5G4^+^ (left) and MJFR14^+^ (right) signals in nucleus, soma, and neurite of TH^+^ neurons in 12M WT, S129A KI, and S129D KI mice (n=6). (J) Multiplex immunofluorescence co-staining (DAPI, TH, 5G4, LAMP1, Ub, TOM20) and 3D-rendered reconstruction of SN sections from 12M S129A KI mice, showing colocalization of 5G4^+^ aggregates with organelle/ubiquitin markers. Scale bar: 5 μm. (K) Multiplex immunofluorescence co-staining (DAPI, TH, MJFR14-6-4-2, TUJ1, Ub, TOM20) and 3D-rendered reconstruction of SN sections from 12M S129A KI mice, showing spatial relationship of MJFR14^+^ aggregates with neurite/organelle/ubiquitin markers. Scale bar: 5 μm. All data are presented as mean ± SEM. Quantification analyses (B, D, F, H) used two-way ANOVA with Tukey’s post hoc test; subcellular localization analysis (I) used one-way ANOVA with Tukey’s post hoc test. α-syn, α-synuclein; KI, knock-in; LAMP1, lysosomal-associated membrane protein 1; PK, proteinase K; SN, substantia nigra; TH, tyrosine hydroxylase; TUJ1, neuron-specific class III β-tubulin; Ub, ubiquitin; TOM20, translocase of outer mitochondrial membrane 20; WT, wild-type.

To confirm the pathological resistance of S129A α-syn aggregates, we performed proteinase K (PK) digestion assays. Without PK treatment, total α-syn positive area was comparable across genotypes at either 3 or 12 M (Fig. 2C, S4A), ruling out an impact of shifting baseline α-syn levels on PK-resistance. After PK treatment, α-syn signals were nearly eliminated across all genotypes at 3 M. By 12 M, however, striking genotype-specific difference emerged: PK abolished α-syn signals in WT and S129D mice, while prominent PK-resistant α-syn^+^ signal persisted in S129A mice (Fig. 2C). Quantification revealed drastically elevated α-syn PK-resistant indices in 12 M S129A mice relative to WT (Fig. 2D). These results indicate that S129A promotes the age-dependent formation of protease-resistant α-syn aggregates.

We further characterized α-syn aggregation via immunofluorescence using aggregation-specific (5G4)(*13*) and conformation-specific (MJFR14-6-4-2)(*14*) antibodies. Antibody specificity was validated in α-syn knockout mice (Fig. S4B). For 5G4, TH co-staining (3–12 M) showed prominent punctate signals in S129A TH^+^ neurons, with minimal fluorescence in WT and S129D mice (Fig. 2E, S4C). Quantification revealed elevated 5G4^+^ areas in S129A TH^+^ neurons as early as 3 M, amplifying further at 6 M and 12 M (Fig. 2F). Western blotting further confirmed 5G4^+^ α-syn enrichment in the insoluble SN fraction of S129A mice at 12 M (Fig. S4E–F). For MJFR14-6-4-2, dense immunoreactivity was detected in S129A TH^+^ neurons, with sparse staining in WT and S129D mice. S129A mice exhibited increased MJFR14^+^ area by 3 M, with elevations by 12 M, whereas S129D mice showed mild elevation only at 12 M (Fig. 2G–H, S4D). Immunohistochemistry further confirmed accumulation of MJFR14^+^ aggregates in the S129A SN (Fig. S4G–H). Notably, 5G4^+^ and MJFR14^+^ aggregates displayed distinct subcellular localizations in 12 M S129A mice: 5G4^+^ aggregates were enriched in TH^+^ neuron soma (secondary nuclear localization), while MJFR14^+^ aggregates were predominantly distributed in neurites (Fig. 2I).

Given that 5G4^+^ aggregates localize to both the soma and nucleus, while MJFR14^+^ aggregates concentrate in neurites, we performed region-specific multiplex co-staining to characterize their molecular associations. For soma-localized 5G4^+^ aggregates, co-staining with lysosomal marker LAMP1, ubiquitin (Ub, a key component of PD pathological inclusions), and mitochondrial marker TOM20 showed clear colocalization(*15*). 3D-rendered images revealed that 5G4^+^ clusters in S129A dopaminergic soma overlapped with LAMP1, Ub, and TOM20 signals—mirroring the organelle-associated, ubiquitinated features of LBs. Nuclear 5G4^+^ aggregates colocalized with Ub, consistent with the ubiquitinated α-syn inclusions reported in PD patients’ neuronal nuclei(*16*) (Fig. 2J). For neurite-localized MJFR14^+^ aggregates, co-staining with neurite marker TUJ1, Ub, and TOM20 demonstrated filament-like signal overlap with Ub and TOM20 (Fig. 2K). 3D-rendering further illustrated MJFR14^+^ aggregates accumulating along TUJ1^+^ neurites with these markers, recapitulating the ubiquitinated, mitochondria-associated properties of Lewy neurites (LNs) in PD.

These results suggest that the compartment-specific aggregates in S129A KI mice mirror some of the hallmarks of PD (LBs and LNs) and their molecular signatures (ubiquitination, mitochondrial association), suggesting that S129A-driven α-syn aggregation recapitulates these PD-relevant pathological processes.

### Mitochondrial, synaptic, and ubiquitin defects in S129A KI mice

To explore the underlying pathogenic mechanisms linking α-syn aggregation and PD-like phenotypes in S129A KI mice, we used spatial transcriptomics (Stereo-seq) to profile molecular features in WT and S129A KI mice (Fig. 3A). Stereo-seq on SN sections generated 32 distinct transcriptomic clusters (Fig. 3B, S5). DA neurons were identified from these clusters via 4 canonical markers: *TH*, *DDC* (dopa decarboxylase), *SLC6A3* (dopamine transporter, DAT), and *SLC18A2* (vesicular monoamine transporter 2, VMAT2); spatial visualization of this population confirmed genotype-specific transcriptomic patterning between WT and S129A (Fig. 3B).

**Fig. 3.**
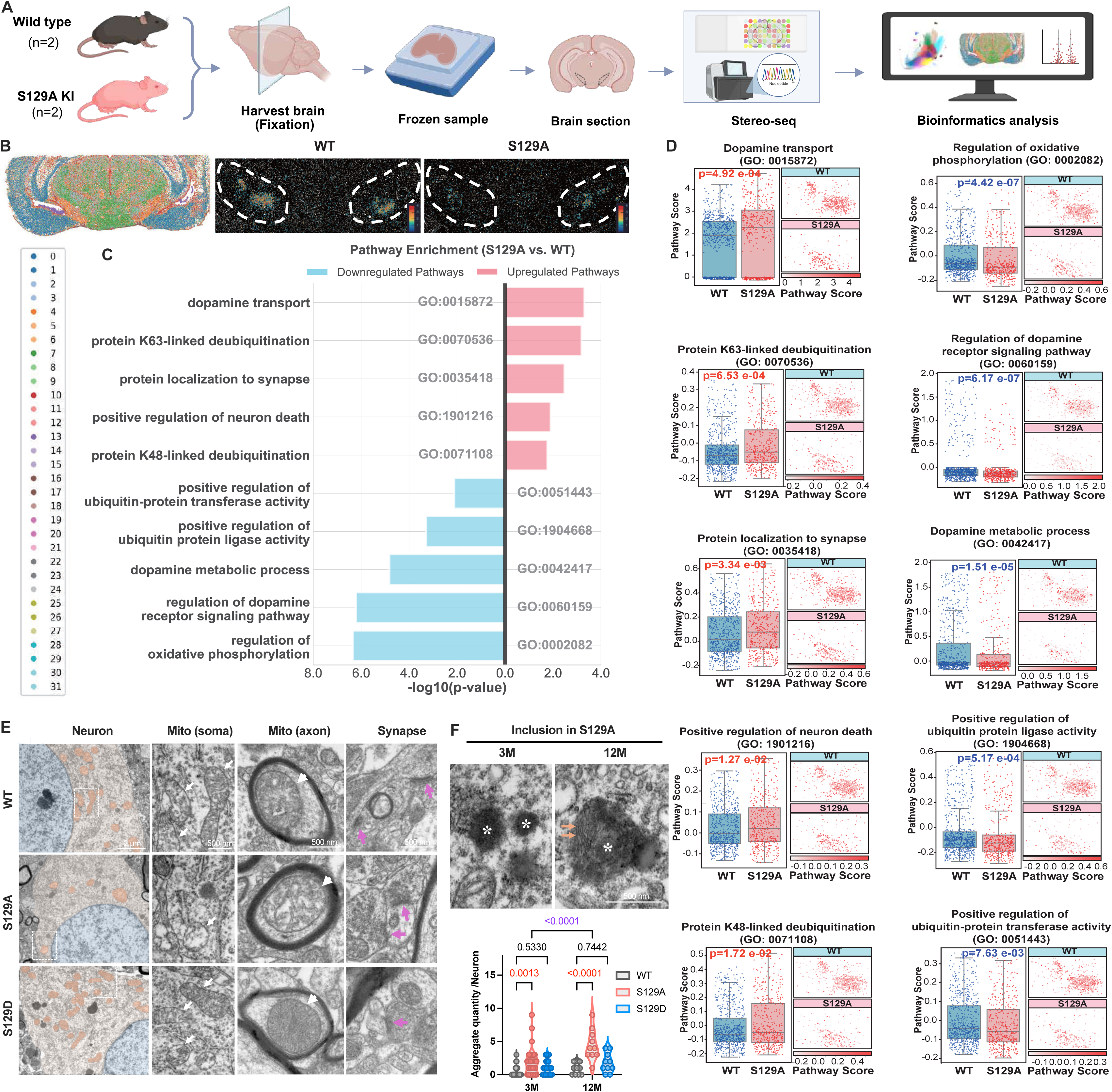
Spatial transcriptomic and ultrastructural profiling identifies PD-relevant pathogenesis in S129A KI mice. (A) Experimental workflow: Brains from WT and S129A KI mice (n=2) were harvested, fixed, processed into FFPE samples, sectioned, analyzed via Stereo-seq for spatial transcriptome profiling, and subjected to bioinformatic analysis. (B) Spatial transcriptome visualization: Left, global map of nigrostriatal sections; right, magnified SN regions showing genotype-specific spatial expression patterning in WT vs. S129A. (C) Pathway scoring analysis (S129A vs. WT): Top 10 dysregulated PD-relevant pathways (ranked by p-value of score differences) are plotted, with upregulated (red) and downregulated (blue) pathways annotated with corresponding GO terms and IDs. (D) Pathway score distributions: Representative plots for dysregulated pathways show pathway score differences between WT and S129A dopaminergic neurons, with statistical significance (p-values) indicated. (E) TEM of SN subcellular structures: Panels display ultrastructure of neurons, somatic mitochondria, axonal mitochondria, and synapses in WT, S129A, and S129D KI mice; white arrows mark mitochondria, purple arrows mark synapses. (F) Pathological aggregates in S129A neurons: Top, TEM images of high-electron-density aggregates (marked by *) at 3 M and 12 M, orange arrows indicate filamentous substructures; bottom, box-and-whisker plot quantifies aggregate quantity per neuron. All data are presented as mean ± SEM. Statistical analysis for pathway score comparisons (D) used Student’s t-test; quantification of aggregate quantity (F) used two-way ANOVA with Tukey’s post hoc test. α-syn, α-synuclein; FFPE, formalin-fixed paraffin-embedded; KI, knock-in; Mito, mitochondria; SN, substantia nigra; TEM, transmission electron microscopy; TH, tyrosine hydroxylase; WT, wild-type.

We extracted DA neurons from both genotypes and scored PD-relevant biological pathways to evaluate their regulatory status. The top 10 significantly dysregulated pathways (Fig. 3C-D) converged on interconnected pathological processes. Mitochondrial function was prominently impaired. Thus, oxidative phosphorylation (GO: 0002082), a core mitochondrial bioenergetic pathway, was markedly downregulated in S129A mice, consistent with the established role of mitochondrial dysfunction in PD-related dopaminergic loss(*17*). Dopamine homeostasis was disrupted: dopamine receptor signaling and metabolism (GO: 0042417) were downregulated, while dopamine transport (GO: 0015872) was upregulated (a potential compensatory response to impaired metabolism). An imbalance between dopamine metabolism and uptake likely perturbs synaptic signaling. Ubiquitin-mediated proteostasis was critically dysregulated: two key ubiquitin-conjugation pathways (positive regulation of ubiquitin protein ligase (GO: 1904668, targeting E3 ligases) and ubiquitin-protein transferase activity (GO: 0051443, governing E2 enzymes) were downregulated, blunting targeted protein ubiquitination. Conversely, K63-linked (GO: 0070536) and K48-linked (GO: 0071108) deubiquitination pathways were upregulated. As K48-linked ubiquitination signals proteasomal degradation(*18*) and K63-linked chains mediate stress response/autophagic processes(*19*), this dysregulation disrupts cellular homeostasis and promotes α-syn aggregation(*20*), key PD mechanisms tied to impaired protein degradation and lysosomal dysfunction(*21*). Synaptic function was also dysregulated: upregulation of protein localization to synapse (GO: 0035418) may compensate for reduced dopamine levels, consistent with the link between α-syn phosphorylation and synaptic function(*7*). Additionally, upregulation of positive regulation of neuron death (GO: 1901216) aligned with S129A-linked TH^+^ neuron loss, reflecting enhanced dopaminergic vulnerability to non-phosphorylated α-syn.

To link these molecular dysregulations to subcellular structures, we performed electron microscopy (EM) on the SN from WT, S129A KI, and S129D KI mice (Fig. 3E). Nuclear integrity was preserved across all genotypes, confirming that subcellular alterations precede cell death. However, mitochondrial morphologies differed markedly, in that WT/S129D neurons had abundant, intact mitochondria with dense cristae (soma/axons), while S129A neurons showed swollen mitochondria, fragmented cristae, and reduced quantities (soma/axons). Synaptic compartments also displayed genotype-specific defects: WT/S129D synapses maintained orderly vesicle pools and distinct pre/post-synaptic membranes, while S129A synapses had disorganized vesicles and blurred synaptic structures. These phenotypes align with the transcriptomic changes, such as compensatory upregulation of synaptic localization pathways (Fig. 3C-D) and downregulation of oxidative phosphorylation, likely reflecting impaired synaptic transmission and neuronal bioenergetics.

Notably, pathological aggregates (Fig. 3F) were exclusive to S129A neurons: high-electron-density aggregates were already present at 3 M, expanding dramatically in size and number by 12 M with filamentous substructures, a signature of α-syn fibrillation. This age-dependent progression directly correlates with ubiquitin-mediated proteostasis dysregulation of S129A (Fig. 3C-D): impaired ubiquitination paired with enhanced deubiquitination may compromise misfolded α-syn clearance, enabling early aggregate onset and subsequent expansion.

Collectively, spatial transcriptomic and EM analyses demonstrate that non-phosphorylated α-syn drives coordinated dysregulation of mitochondrial, proteostasis, and dopamine-synaptic pathways, alongside subcellular defects, consistent with canonical hallmarks of PD pathology. This reinforces the notion that S129A KI mice recapitulate PD-relevant pathogenic features, such as the association with mitochondrial components, aligning with human proteomic data on non-phosphorylated human α-syn(*22*).

### Temporally stratified non-motor symptoms in S129A KI mice

PD features not only motor deficits but also temporally stratified non-motor symptoms across disease stages: olfactory dysfunction and gastrointestinal impairments typically emerge during the prodromal phase, clinically preceding motor symptoms by up to 20 years, while cognitive decline manifests in later stages, after motor symptom onset (*23*). To determine whether S129A KI mice recapitulate this clinical progression, we systematically assessed olfactory, gastrointestinal, and cognitive phenotypes across different ages.

To evaluate olfactory function, we performed the buried food and odor preference tests (Fig. 4A). As early as 3 M, S129A mice exhibited significantly prolonged latencies to locate buried food vs. WT, with this deficit persisting through 6, 9, 12 M (Fig. 4B); S129D mice showed no differences. All genotypes had comparable surface food-seeking latencies (Fig. S6A), ruling out non-olfactory factors (e.g., motor/emotional impairments). We further validated this with the odor preference task, which assesses responses to attractive (2-phenylethanol), neutral (isoamyl acetate), and aversive (2,4,5-trimethylthiazole) odors(*24*): S129A mice displayed reduced preference for attractive and aversive odors as early as 3 M compared to WT mice, while S129D mice also showed abnormal alterations in responses to attractive odors (Fig. 4C). For structural correlates, Nissl staining of the olfactory bulb (OB) focused on the mitral cell layer (MCL), which is critical for olfactory signal transmission(*25*). S129A mice had significantly reduced MCL Nissl^+^ cell density at 3 M vs. WT, while S129D mice showed no differences (Fig. 4D–E). Analysis of all OB layers (GCL, IPL, MCL, EPL, GL) revealed S129A mice had reduced Nissl^+^ cell densities across the total layered structure of the OB, while S129D mice showed milder, layer-specific (GCL, GL) reductions (Fig. S6B–C). These results demonstrate that early olfactory dysfunction in S129A mice is accompanied by pan-OB structural deficits, with S129D mice exhibiting milder, layer-specific alterations.

**Fig. 4.**
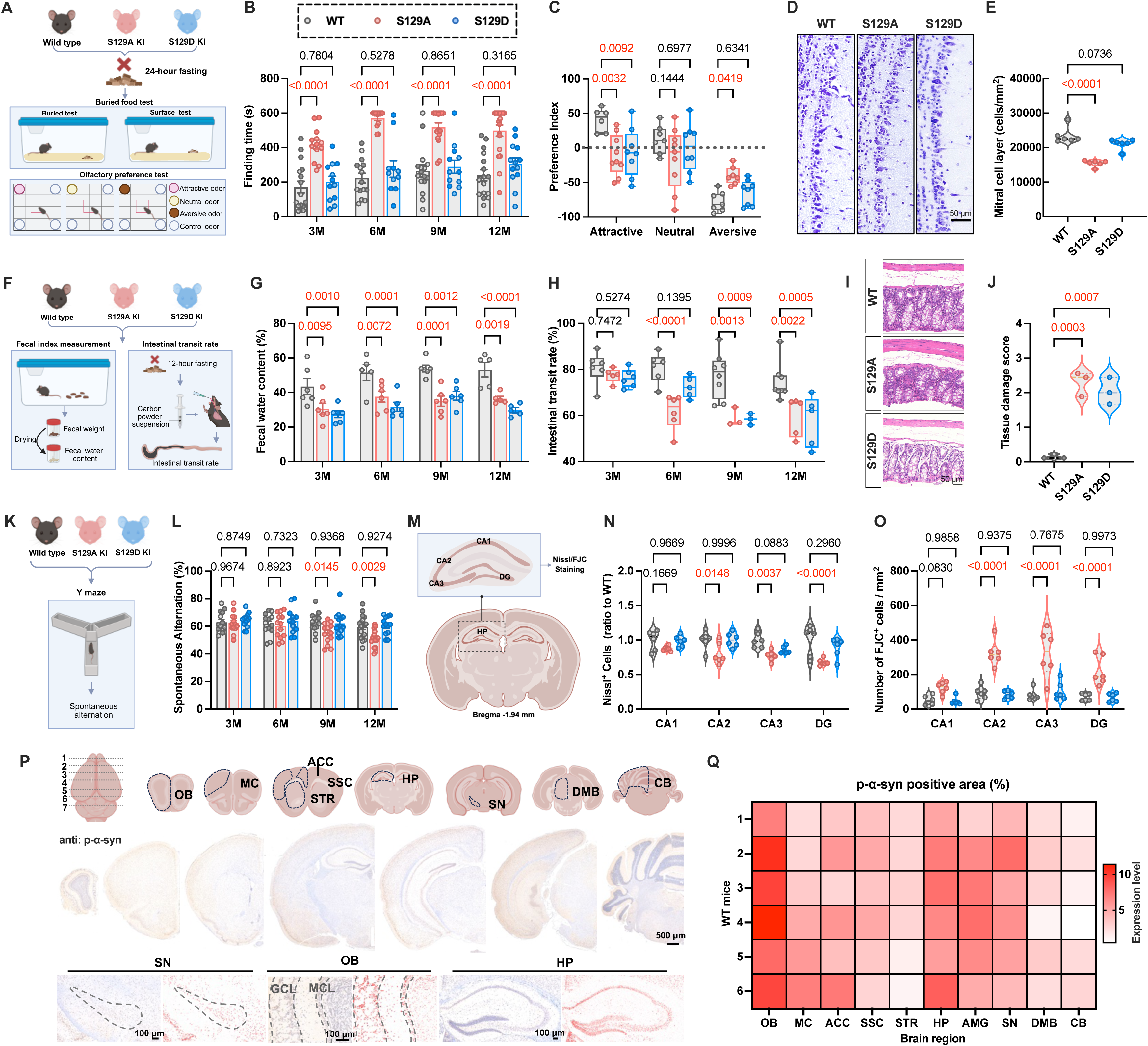
S129A knock-in mice recapitulate temporally stratified PD-like non-motor symptoms. (A) Schematic of experimental designs for assessing olfactory function: buried/surface food tests and odor preference test conducted in WT, S129A KI, and S129D KI mice. (B) Latency to locate food in the buried food tests across genotypes and ages (n=11-19, including males with dark points and females with light points). (C) Odor preference index for attractive (2-phenylethanol), neutral (isoamyl acetate), aversive (2,4,5-trimethylthiazole) odors across genotypes (n=6-9). (D) Nissl staining of the olfactory bulb (scale bar: 50 μm) across genotypes. (E) Quantification of Nissl^+^ cell density in the MCL of the OB (n=6). (F) Schematic of experimental designs for assessing gastrointestinal function: fecal water content and intestinal transit rate assessment. (G) Fecal water content across genotypes and ages (n=5-6). (H) Intestinal transit rate across genotypes and ages (n=3-8). (I) Histological staining of colon tissue (scale bar: 50 μm) across genotypes. (J) Quantification of colon tissue damage score (n=3). (K) Schematic of the Y-maze test for assessing working memory. (L) Spontaneous alternation rate in the Y-maze test across genotypes and ages (n=13-16, including males with dark points and females with light points). (M) Schematic of the HP indicating subregions (CA1, CA2, CA3, DG) assessed via Nissl/FJC staining. (N) Quantification of Nissl^+^ cell density (normalized to WT) in hippocampal subregions (n=6). (O) Quantification of FJC^+^ degenerating neuron number in hippocampal subregions (n=6). (P) p-α-syn immunohistochemistry of multiple WT mouse brain regions (including OB, MC, ACC, SSC, STR, HP, SN, DMB, CB; scale bars: 500 μm and 100 μm).(Q) Heatmap of p-α-syn positive area percentage across WT mouse brain regions (n=6). All data are presented as mean ± SEM. Analyses for B, C, G, H, L, N, O used two-way ANOVA with Tukey’s post hoc test; analyses for E and J used one-way ANOVA with Tukey’s post hoc test. ACC, anterior cingulate cortex; CB, cerebellum; DMB, dorso-medial midbrain; DG, dentate gyrus; FJC, Fluoro-Jade C; HP, hippocampus; KI, knock-in; MC, motor cortex; OB, olfactory bulb; SSC, somatosensory cortex; SN, substantia nigra; STR, striatum; WT, wild-type.

For gastrointestinal function assessment, we evaluated constipation tendency, via fecal water contents and intestinal transit rates (Fig. 4F). S129A and S129D mice had reduced fecal weights compared with WT mice across ages (Fig. S6D). Both genotypes showed significantly lower fecal water content compared with WT from 3 M onward (Fig. 4G), consistent with prodromal constipation, a characteristic PD symptom that precedes motor deficits by decades. S129A mice had impaired intestinal transit by 6 M (early-onset motility dysfunction), while S129D mice showed comparable deficits with delayed onset (9 M; Fig. 4H). Colon histology revealed S129A mice had increased outer longitudinal muscle layer thickness; both S129A/S129D mice exhibited tissue damage and immune cell infiltration compared with WT (Fig. 4I–J, S6E–F). Tight junction protein (ZO-1, Claudin-5) immunofluorescence confirmed reduced colonic barrier integrity in both S129A/S129D mice (Fig. S6G–I). Open field tests showed no gross locomotor differences (distance, time, speed) across genotypes/ages (Fig. S6J–L), ruling out locomotor effects on gastrointestinal deficits. Collectively, S129A mice recapitulated prodromal gastrointestinal symptoms, aligning with other PD-relevant phenotypes. S129D mice also exhibit gastrointestinal abnormalities—hinting that the function of α-syn S129 phosphorylation differs between the gastrointestinal tract and CNS, with its regulatory mechanism requiring further investigation.

To evaluate cognitive function (a late PD symptom post-motor onset), we used the Y-maze test to assess spatial working memory (Fig. 4K). S129A mice exhibited significantly lower spontaneous alternation rates compared with WT starting at 9 M, with further worsening by 12 M—notably later than their motor symptom onset (Fig. 4L), mirroring clinical late-stage cognitive impairments in most subjects with PD. To explore the structural basis, we performed Nissl (neuron density) and FluoroJade-C (FJC; degenerating neuron) staining of the hippocampus at 12 M (Fig. 4M). S129A mice had reduced Nissl^+^ cell density across all hippocampal subregions (CA1, CA2, CA3, DG) compared with WT (Fig. 4N, S6M), alongside markedly increased FJC^+^ degenerating neurons (Fig. 4O, S6N), suggesting that hippocampal neurodegeneration underlies their cognitive deficits. In contrast, S129D mice showed no significant Y-maze cognitive impairment, nor hippocampal neuronal loss or degeneration.

Given that S129A KI mice exhibit PD-like symptoms and multi-regional brain damage, we suspected that endogenous pS129 α-syn might have a physiological role in these vulnerable regions, based on multiple recent reports(*7, 8, 26, 27*)—although it was traditionally regarded as a pathological marker because it was suggested to account for <4% of total α-syn globally(*5*). Thus, we measured pS129 expression via immunohistochemistry on WT mouse brains, covering regions from the olfactory bulb (OB) to cerebellum (CB): OB, motor cortex (MC), anterior cingulate cortex (ACC), somatosensory cortex (SSC), striatum (STR), hippocampus (HP), substantia nigra (SN), extranigral or dorsomedial midbrain (DMB), and CB (Fig. 4P). No pS129 signal was detected in α-syn knockout (KO) mice (Fig. S7A), suggesting antibody specificity. Quantitative analysis revealed striking regional differences: SN, OB, and HP showed markedly high p-α-syn positive area ratios (Fig. 4P–Q), consistent with prior reports of region-specific high α-syn expression in WT mice (*28*). Further immunofluorescence co-staining confirmed pS129 α-syn colocalized with TH^+^ dopaminergic neurons in WT SNpc (Fig. S7B), while no signal was detected in S129A or S129D KI SNpc due to the missing phospho-serine residue (Fig. S7C). This observation may help resolve prior misperceptions: pS129 α-syn is not inherently pathological but exhibits region-specific high expression in PD-vulnerable brain regions, precisely in those areas where S129A mice show pathological damage. These locally high levels likely support regional physiological functions that are disrupted by loss of S129 phosphorylation in S129A mice, aligning with our suspicions and directly explaining their PD-like phenotypes.

In summary, relative to other traditional genetic models(*29, 30*), the S129A KI mice appear to better recapitulate the temporally stratified non-motor symptoms of PD, such as prodromal olfactory and gastrointestinal deficits, followed by late-stage cognitive impairment. The discovery of region-specific high endogenous pS129 α-syn expression further supports its physiological role, reinforcing that S129A KI mice mirror PD’s clinical progression and pathogenesis linked to impaired α-syn S129 phosphorylation.

### Amelioration of early S129A phenotypes via phosphorylation-restoring/aggregation-targeting interventions

Leveraging our new findings that S129A KI mice develop PD-like symptoms tied to impaired α-syn S129 phosphorylation, and that endogenously pS129 α-syn is regionally enriched in PD-vulnerable brain regions, we hypothesized that modulating α-syn phosphorylation might alleviate these preclinical phenotypes. We first evaluated whether partial restoration of endogenous α-syn phosphorylation could mitigate the deficits. We analyzed S129A heterozygous mice (S129A^+^/^−^), which retain partial capacity for endogenous α-syn phosphorylation (unlike S129A homozygotes, S129A^+^/^+^) (Fig. 5A). At 3 M, when S129A^+^/^+^ mice already exhibit olfactory and gastrointestinal deficits, S129A^+^/^−^ mice showed a significantly improved relative change rate of food-finding latency (Fig. 5B), alongside rescued fecal weight and fecal water content (Fig. 5C–D). These results suggested that even partial restoration of α-syn phosphorylation mitigates early S129A-associated disease phenotypes.

**Fig. 5.**
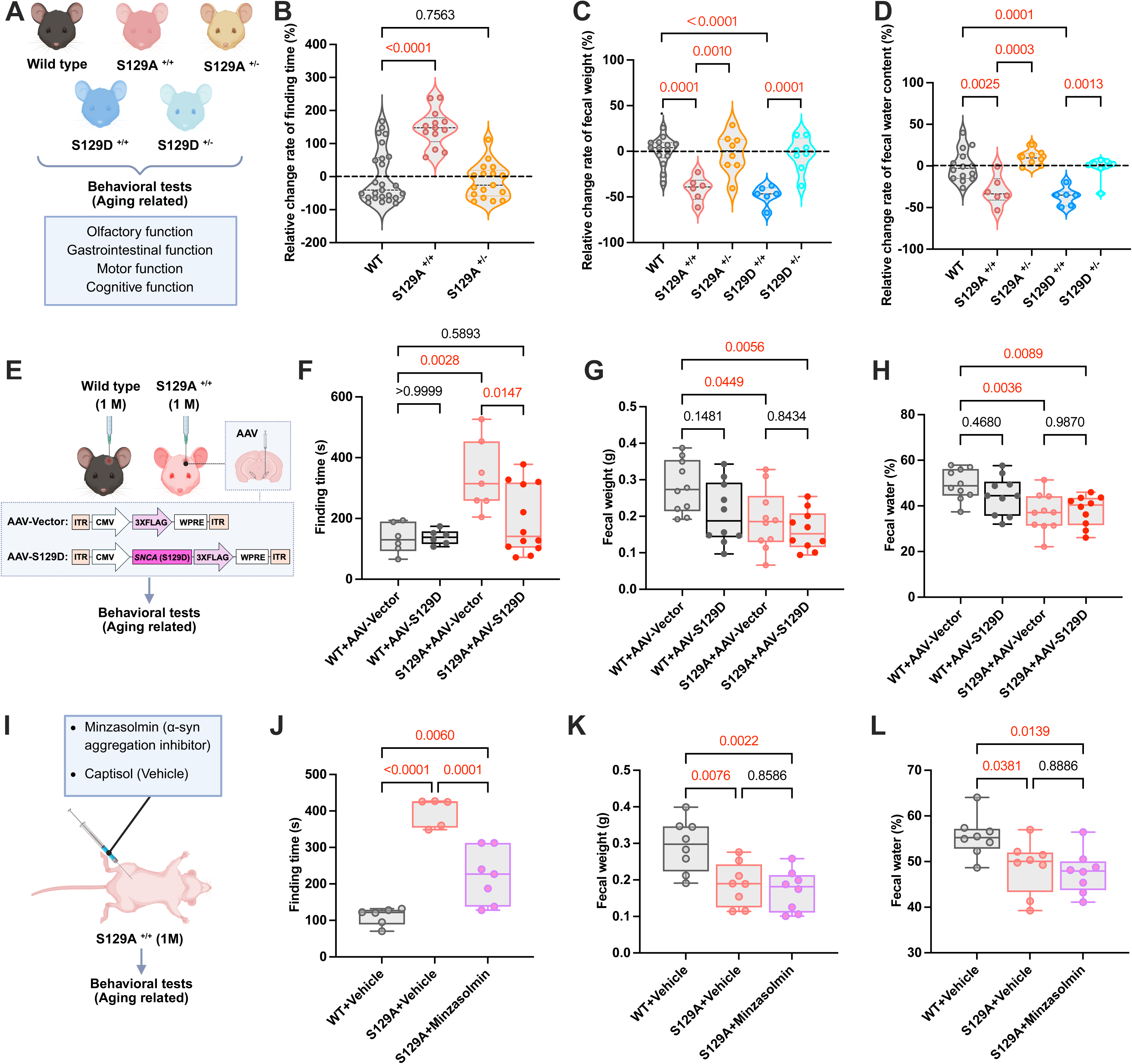
Restoring S129 phosphorylation or targeting α-syn aggregation ameliorates prodromal symptoms in S129A knock-in mice. (A) Schematic of experimental design for assessing aging-related symptoms in WT, S129A homozygous (S129A^+^/^+^), and heterozygous (S129A^+^/^−^) mice. (B) Relative change in food-finding latency across genotypes (n=13-25). (C) Relative change in fecal weight across genotypes (n=6-14). (D) Relative change in fecal water content across genotypes (n=6-14). (E) 1-month-old WT and S129A^+^/^+^ mice injected with AAV-S129D (to restore S129 phosphorylation) or AAV-Vector (control), followed by aging-related behavioral assessments. (F) Food-finding latency in AAV-treated mice (n=6-12). (G) Fecal weight in AAV-treated mice (n=10). (H) Fecal water content in AAV-treated mice (n=10). (I) 1-month-old S129A^+^/^+^ mice administered Minzasolmin (α-syn aggregation inhibitor) or Captisol (vehicle), followed by aging-related behavioral assessments. (J) Food-finding latency in Minzasolmin/vehicle-treated S129A^+^/^+^ mice (n=per group sample size). (K) Fecal weight in Minzasolmin/vehicle-treated S129A^+^/^+^ mice (n=5-7). (L) Fecal water content in Minzasolmin/vehicle-treated S129A^+^/^+^ mice (n=8). All data are presented as mean ± SEM. Statistical analyses used one-way ANOVA with Tukey’s post hoc test. AAV, adeno-associated virus; KI, knock-in; WT, wild-type.

To confirm that restoring α-syn phosphorylation directly drives this amelioration, we used a targeted, region-specific intervention. We delivered AAV-driven S129D to the SN of S129A^+^/^+^ mice starting at 1 M—before the onset of detectable pathological changes (Fig. 5E). This midbrain region is not only a core PD-vulnerable area but also one where endogenous pS129 α-syn is highly expressed in WT mice. At 3 M, S129A^+^/^+^ mice receiving AAV-S129D, but not empty AAV vector, exhibited significantly shorter food-finding latency (Fig. 5F), indicating hyposmia amelioration. Notably, fecal weight and water content remained unaltered relative to S129A^+^/^+^ mice injected with empty AAV vector (Fig. 5G–H). This selective improvement likely reflects nigral regulation of olfactory function, mediated by neural projection pathways linking the medioventral SN to olfactory processing circuits(*31, 32*), whereas gastrointestinal homeostasis may depend on distinct α-syn-expressing networks or require more widespread restoration of S129 phosphorylation beyond the SN. Collectively, these data reinforce the notion that restoring α-syn phosphorylation in key PD-vulnerable brain regions alleviates clinically relevant early S129A phenotypes.

Beyond phosphorylation restoration, we targeted α-syn aggregation—another key pathological feature that dovetails with our prior observations of dopaminergic neuronal damage in S129A mice. We administered Minzasolmin, an α-syn aggregation inhibitor(*33*), to S129A^+/+^ mice starting at 1 M (Fig. 5I). At 3 M, Minzasolmin treatment improved food-finding latency (Fig. 5J), confirming that targeting α-syn aggregation is an effective approach to alleviate olfactory dysfunction in S129A mice. Interestingly, unlike the partial restoration of phosphorylation in heterozygous mice, Minzasolmin did not rescue fecal weight or water content (Fig. 5K–L). This outcome highlights that gastrointestinal function may be regulated by more intricate mechanisms, representing another important direction for future exploration.

Notably, motor and cognitive functions remained comparable across all groups in the three aforementioned interventions at 3 M (Fig. S8–S10), given that the mice had not yet progressed to the stages of motor or cognitive symptom onset. We are continuing to monitor these phenotypes to evaluate long-term interventional effects in aging mice.

In summary, S129A KI mice recapitulate key manifestations of PD, with temporally stratified phenotypes and LB-like aggregates. Notably, both phosphorylation-modulating and aggregation-targeting treatments alleviate PD-related deficits in S129A KI mice, confirming the pathogenicity of α-syn S129 phosphorylation loss and reinforcing the translational value of S129A KI.

## Discussion

This study capitalizes on two genetically precise KI models to address a core controversy—whether p-α-syn is pathogenic or protective—to reshape several questions in the PD field. S129A mice recapitulate key aspects of the clinical trajectory of PD, while S129D mice phenocopied WT controls in all but the GI measures, alongside mild olfactory deficits. These new observations support the view that *loss* of S129 phosphorylation triggers PD progression, rather than the decades-old dogma that p-α-syn drives PD pathogenesis (Fig. S11).

Existing PD models (transgenic, neurotoxic, preformed fibrils, etc.) have made invaluable contributions to elucidating pathogenic features of PD(*29*), such as dopaminergic neuron loss. However, the field has long sought a genetic model that better mirrors the natural, temporally stratified trajectory of human PD. We believe that the S129A model recapitulates this temporal progression of clinical features more closely than traditional murine models. Thus, the S129A model induces prodromal olfactory/gastrointestinal deficits at 3 months, progressive motor dysfunction at 6 months, and cognitive decline at 9 months, with each stage accompanied by PD-relevant histopathological and biochemical changes. Critically, the temporal alignment with clinical non-motor symptoms preceding motor deficits delivers a relatively unique *in vivo* tool for dissecting disease mechanisms and testing stage-specific interventions. These important new features may reorient therapeutic strategies.

The validation of the S129A model helps address a longstanding controversy surrounding p-α-syn. Long viewed as a pathogenic component of LBs, p-α-syn has steered most therapeutic efforts toward *inhibiting* S129 phosphorylation(*5, 34*). Our data may resolve this ambiguity: a lack of α-syn phosphorylation appears to initiate PD-like pathogenesis, and interventional data further suggest that restoring S129 phosphorylation mitigates preclinical neurological symptoms. This reversal not only addresses a decade-long debate but also shifts the therapeutic paradigm from inhibiting S129 phosphorylation to preservation of its physiological roles. Second, it raises the possibility that environmental disease triggers may initiate non-familial PD by inhibiting the physiological phosphorylation of α-syn at this residue, perhaps encouraging aggregate formation and spread from neurons that interface with the gut and nasal mucosae.

Building on these findings, multiple directions merit further exploration. For humanized models, generating human S129A KI models, patient-derived iPSC lineages(*35*), and non-human primate S129A models(*36, 37*) will advance mechanistic research and therapeutic development. For prodromal mechanisms, the early olfactory/gastrointestinal deficits of the S129A model improve our understanding of disease initiation mechanisms and help identify earlier intervention time windows(*23*). For therapeutic optimization, modulating S129 phosphorylation emerges as a key strategy(*38*), and yet, its efficacy may be shaped by heterogeneity across tissues, cell types, and disease stages. This heterogeneity warrants in-depth exploration to guide combinatorial/stage-specific therapies.

## Materials and Methods

### Mice, Study Design, and Ethics

All animal procedures were approved by the Animal Care and Use Committee of Capital Medical University and complied with national guidelines for the care and use of laboratory animals (AEEI-2025-1094). All knock-in lines were maintained on a C57BL/6J background. S129A and S129D Snca knock-in mice were generated by CRISPR/Cas9-mediated point mutation of Ser129 in exon 5 of the endogenous *Snca* gene, as detailed in “Generation and Genotyping of S129A and S129D Knock-in Mice.” Human α-syn transgenic mice (C57BL/6N-Tg(Thy1-SNCA)15Mjff/J) (Strain #017682) and *Snca* knockout mice (Strain #016123) were purchased from The Jackson Laboratory and bred in-house. These lines were used in select experiments as positive or negative controls for α-syn and p-α-syn detection.

All mice were housed under specific pathogen–free conditions on a 12 h light–dark cycle with controlled temperature and humidity, and had ad libitum access to food and water. Both male and female mice were used in this study. Unless otherwise stated, age-matched animals from the same colony were used as controls. Mice were randomly assigned to genotype and treatment groups, and data collection and analysis were performed with the experimenter blinded to genotype and treatment whenever feasible.

### Generation and Genotyping of S129A and S129D Knock-in Mice

S129A and S129D knock-in mice were generated in 2021 on a C57BL/6J background using the CRISPR/Cas9 technology (Shanghai Model Organisms, Shanghai, China). CRISPR/Cas9-mediated homology-directed repair targeted exon 5 of the endogenous mouse *Snca* gene (Ensembl gene ID: ENSMUSG00000025889; transcript Snca-201, ENSMUST00000114268.5). In both lines, a single guide RNA (sgRNA) recognizing exon 5 (5′-GGCTTATGAAATGCCTTCAG-3′; PAM: AGG) was used together with a mutation-specific single-stranded oligodeoxynucleotide (ssODN) donor template. For the S129A line, the codon encoding Ser129 (TCA) was replaced by GCA; for the S129D line, the same Ser129 codon was replaced by GAC. The ssODN donors additionally contained synonymous substitutions within the sgRNA/PAM region to prevent re-cutting by Cas9 and to facilitate genotyping.

Cas9 mRNA, sgRNA, and ssODN donors were synthesized in vitro and co-injected into fertilized C57BL/6J zygotes using standard pronuclear microinjection procedures. Injected embryos were transferred into pseudopregnant recipient females, and F0 offspring were screened for the desired point mutations by PCR amplification of the targeted exon 5 region followed by Sanger sequencing of the PCR products. Mosaic founders carrying the correct S129A or S129D substitutions without undesired insertions or deletions were crossed with C57BL/6J mice to generate F1 heterozygous carriers. Heterozygous offspring were then intercrossed to obtain the wild-type, heterozygous, and homozygous S129A or S129D knock-in mice used in this study.

For routine genotyping, tail genomic DNA was amplified using primer pairs flanking the edited region. For the S129A line, PCR was performed with P1 (5′-ACTTCGTGCAGCACCTTGAA-3′) and P2 (5′-TGCCCGTTGTTACTCAGACA-3′). For the S129D line, PCR primers were P1 (5′-AACACTTCGTGCAGCACCTT-3′) and P2 (5′-ACTGCCCGTTGTTACTCAGA-3′). Genotypes (wild-type, heterozygous, or homozygous knock-in) were determined by direct sequencing of PCR products and inspection of the chromatograms at the edited codon. Experimental cohorts consisted of age- and sex-matched wild-type and homozygous knock-in mice derived from the same colony. Full sequences of the wild-type locus, ssODN donors, sgRNA, and genotyping primers are provided in Supplementary Table S2.

### Adeno-Associated Virus (AAV) Vector Construction and Stereotaxic Injection

Recombinant adeno-associated virus serotype 8 (AAV8) vectors expressing human *SNCA* carrying the S129D mutation (AAV8–CMV–*SNCA*(S129D)–3Flag) or a corresponding control vector were generated using the GV829 backbone (CMV–MCS–3×Flag–WPRE–BGH polyA). Human *SNCA* cDNA (S129D point mutation) was cloned into the MCS downstream of the CMV promoter and in-frame with a C-terminal 3×Flag tag. Viral packaging and purification were performed by a commercial provider according to standard AAV8 production procedures (GeneChem, Shanghai, China). Final viral preparations were supplied at a titer of 5 × 10^12 vector genomes (vg)/mL and stored at −80 °C until use.

For unilateral nigral injections, 1-month-old WT and S129A mice received AAV8–*SNCA*(S129D)–3Flag or control AAV8 into the left substantia nigra. Mice were anesthetized and placed in a stereotaxic frame, and a small burr hole was made above the target site. A total volume of 1 μL of viral suspension (5 × 10^12 vg/mL) was injected into the left substantia nigra at the following coordinates relative to bregma: anteroposterior (AP) −2.7 mm, mediolateral (ML) +0.85 mm, dorsoventral (DV) −3.3 mm, based on a previously reported protocol for nigral AAV delivery(*39*). Virus was delivered slowly over several minutes, after which the needle was left in place briefly before being withdrawn to minimize reflux. The scalp incision was closed, and mice were placed in a warmed recovery cage and monitored until fully ambulatory.

### Minzasolmin Treatment

Minzasolmin is a small-molecule inhibitor of α-syn aggregation. In this study, it was administered chronically in vivo to suppress pathological α-syn. Minzasolmin (UCB0599, MedChemExpress, NJ, USA) was dissolved in a vehicle solution containing 40% Captisol (Aladdin, Shanghai, China) in sterile water. The drug or vehicle was injected intraperitoneally at a volume of 0.1 mL per 20 g body weight.

Mice were assigned to vehicle or Minzasolmin treatment groups at 1 month of age. Animals received one intraperitoneal injection per day from Monday to Friday (1 mg/kg Minzasolmin or vehicle) for approximately 60 days, until they reached 3 months of age, following a regimen adapted from a previous study(*33*). After completion of the dosing period, mice underwent behavioral testing, followed by tissue collection for neuropathological and biochemical analyses.

All solutions were prepared and coded by an experimenter who was not involved in data acquisition. Investigators performing behavioral testing, tissue processing, imaging, and quantitative analyses were blinded to genotype and treatment until all data were processed and predefined exclusion criteria were applied.

### Behavioral Tests

#### Rotarod Test

Motor coordination and motor learning were assessed using an accelerating rotarod (Rota Rod System, Panlab, Harvard Apparatus, model 76–0770). Mice underwent a 5-day training period followed by a testing session. During training, animals received one 5-min trial per day. On the first 2 days, an acclimation paradigm was used in which the rod accelerated from 0 to 20 rpm over 300 s. On days 3–5, the training paradigm matched the test condition, with the rod accelerating from 0 to 40 rpm over 300 s. For the test, mice were placed on the rotating drum, and the speed was linearly increased from 4 to 40 rpm within 5 min(*40*). The latency to fall, defined as the time until the mouse either fell from the rod or made a full passive rotation without walking, was recorded over three trials and averaged for each animal. The rod and surrounding compartments were wiped with 75% ethanol between trials.

#### Pole Test

The pole test was used to assess motor coordination and bradykinesia. Mice were tested on a vertical metal pole (48 cm in height, 12 mm in diameter) fixed to a stable base. The top of the pole was covered with a solid sponge ball (∼2 cm in diameter), and the surface of the rod was wrapped with cloth tape to provide grip and prevent slipping. Mice were placed head-up on the sponge ball at the top of the pole, and the time required to descend from the top to the base of the pole was recorded(*41*). Before testing, animals were habituated to the apparatus over two consecutive days, with three training trials per day. On the third day, each mouse performed three test trials, and the mean descent time was used for analysis.

#### Open Field Test

The open field test was used to assess spontaneous locomotor activity and exploratory behavior. The apparatus consisted of a square box (40 × 40 × 40 cm, width × length × height), with a central 20 × 20 cm zone defined by a video tracking system (SMART 3.0, Panlab, Harvard Apparatus). Mice were habituated to the open field for three consecutive days (one 5-min session per day). On the test day, each mouse was placed gently in a corner of the arena and allowed to freely explore for 5 min while behavior was recorded by an overhead camera. SMART 3.0 software was used to quantify total distance traveled, mean locomotor speed, number of entries into the central zone, and the ratio of time or distance in the center relative to the total arena. The arena was thoroughly cleaned with 75% ethanol between trials.

#### Buried Food Test

Olfactory function was assessed using the buried food test, which measures the latency to locate a hidden food pellet. Mice were first habituated to a clean cage containing ∼3 cm of bedding for 15 min per day on two consecutive days. At the end of the second day, food was removed, and animals were food-deprived for 24 h with free access to water. On the test day, a single food pellet was buried approximately 0.5 cm beneath the bedding surface near one corner of the cage. Each mouse was placed in the center of the cage, and the trial was started immediately. The latency to uncover and begin eating the pellet was recorded for up to 10 min using an overhead video tracking system(*42*). Mice that failed to find the food within 10 min were assigned a latency of 600 s for analysis.

To control for basal motivation and food-seeking ability, a visible food test was performed on the following day. Mice were placed in a cage with an unburied food pellet, and the latency to approach and eat the pellet was measured under the same recording conditions.

#### Olfactory Preference Test

Olfactory-driven exploration was assessed in a square arena (40 × 40 × 40 cm, width × length × height) that was virtually divided into four equal quadrants by the video-tracking system. Each mouse was first allowed to freely explore the arena for 10 min to habituate.

On odor trials, a 4 × 4 cm piece of filter paper impregnated with 1 μL of a pure odorant (2-phenylethanol [2-PE], 2,4,5-trimethylthiazole [nTMT], or isoamyl acetate) was placed on the floor in one quadrant(*24*). The mouse was placed in the center of the arena and allowed to explore for 3 min. Sniffing was defined as the time during which the animal’s snout was maintained within ∼2 cm of the filter paper. Time spent in each quadrant and time spent sniffing the odor-containing filter paper were recorded.

Odor preference was quantified using a performance index (PI) that reflects deviation from chance occupancy of the odor quadrant:

PI = (P − 25) / 0.25, where P is the percentage of total time spent in the odor quadrant (chance level = 25%)(*43*) To control for spatial bias, mice were first tested in a 10-min no-odor session; if an animal avoided any quadrant by more than 20% from chance occupancy, the session was discarded and the mouse was retested on another day.

#### Feed intake

Food intake was monitored in group-housed mice with ad libitum access to water and chow. Measurements were collected over 5 consecutive days. On day 1, the total body weight per cage (sum of individual body weights) and the amount of chow present in the hopper were recorded. Each subsequent day at the same time, the chow remaining in the hopper was weighed. Food intake for each 24-hour interval was calculated per cage as the decrease in chow weight between two consecutive daily measurements (day 1–2, day 2–3, day 3–4, and day 4–5), generating one intake value per cage per interval. On day 5, the total body weight per cage was recorded again. Intake values were normalized to body weight (Feed/BW) using the mean of the total cage body weight measured on days 1 and 5.

#### Fecal Pellet Output and Water Content

Gastrointestinal motility and stool water content were assessed by monitoring fecal pellet output and calculating fecal water content over a 120-min period. Mice were fasted overnight in clean cages without bedding, with free access to water. At the start of the experiment, water was removed and each mouse was transferred to an individual cage and observed for 120 min.

Fecal pellets were collected every 30 min immediately after expulsion and placed into pre-weighed, sealed 1.5 mL tubes to minimize evaporation. Tubes were weighed to obtain the total wet weight of feces (A). The samples were then dried in an oven for 24 h and reweighed to obtain the dry weight (B) (*44*). Fecal water content (WCF) was calculated as:

WCF (%) = (A − B) / A × 100%.

#### Small Intestinal Transit Rate

Small intestinal transit was assessed using a carbon meal. The gavage solution contained 4 g sucrose, 8 g milk powder, 1.5 g activated carbon powder, and 5 g sodium carboxymethyl cellulose (CMC-Na) dissolved in 150 mL deionized water. Reagents were added sequentially to a 200 mL volumetric flask, with CMC-Na added in five portions (1 g each) and mixed on a magnetic stirrer until a homogeneous viscous black–gray suspension was obtained.

Each mouse received 150 μL of the carbon meal by oral gavage using a 21-gauge round-tip feeding needle. Thirty minutes after administration, mice were sacrificed and the entire small intestine was carefully dissected from the pylorus to the ileocecal junction. The total length of the small intestine and the distance traveled by the leading edge of the carbon meal were measured and documented(*45*). Small intestinal transit was calculated as:

Small intestinal transit (%) = (distance traveled by carbon meal / total small intestinal length) × 100.

#### Y-Maze Spontaneous Alternation

Working memory was assessed using the spontaneous alternation behavior (SAB) paradigm in a Y-maze. The apparatus consisted of three identical arms (40 × 4 × 9.5 cm) arranged at 120° from each other and designated as arms A, B, and C. Mouse behavior was recorded and analyzed using a video tracking system(*46*).

At the start of each trial, a mouse was placed at the center of the maze and allowed to freely explore all three arms for 8 min. An arm entry was defined as all four paws entering an arm. Spontaneous alternation behavior was calculated as the percentage of triads in which the mouse entered three different arms on consecutive choices (e.g., ABC, BCA, CAB), using the formula: SAB (%) = (number of actual alternations / number of alternation opportunities) × 100, where the number of alternation opportunities equals the total number of arm entries minus 2.

#### Perfusion, Tissue Collection, and Processing

For histological and immunohistochemical analyses, mice were deeply anesthetized and transcardially perfused with ice-cold 0.01 M PBS followed by 4% paraformaldehyde (PFA) in 0.1 M phosphate buffer. Brains were carefully removed from the skull, post-fixed in 4% PFA at 4 °C for 12–24 h, and then processed for paraffin embedding or cryosectioning as described in the corresponding sections. Midbrain blocks containing the substantia nigra and ventral tegmental area, as well as other regions of interest, were dissected according to stereotaxic coordinates and processed in parallel across genotypes and treatment groups.

For biochemical assays, including high-performance liquid chromatography (HPLC) for dopamine and metabolites, sequential protein extraction, and western blotting, mice were sacrificed by rapid decapitation without PFA perfusion. Brains were rapidly removed, and the striatum, midbrain, or other regions of interest were dissected on ice, snap-frozen in liquid nitrogen, and stored at −80 °C until use. For spatial transcriptomics, fresh brains were embedded in OCT and frozen, and coronal sections were collected onto Stereo-seq chips as described above. Peripheral organs (colon, heart, liver, spleen, lung, and kidney) were collected for hematoxylin–eosin staining either after PBS/PFA perfusion or rapid PBS perfusion, fixed in 4% PFA, and processed for paraffin embedding.

For transmission electron microscopy, small tissue blocks (approximately 1 mm³) containing the substantia nigra were dissected immediately after perfusion fixation and further post-fixed in glutaraldehyde, osmicated, dehydrated, and embedded in epoxy resin for ultrathin sectioning.

#### Immunohistochemistry (IHC)

To detect aggregated α-syn in the mouse midbrain, immunohistochemistry was performed using the conformation-specific antibody MJFR14-6-4-2. For immunostaining, sections were deparaffinized in xylene, rehydrated through graded ethanol, and rinsed in distilled water. Antigen retrieval was carried out in 10 mM sodium citrate buffer (pH 6.0) at 85–95 °C for 20–30 min in a water bath, followed by cooling the sections to room temperature. Endogenous peroxidase activity was quenched with 0.3% H O in methanol for 10 min. After washing in PBS, sections were blocked in 5% bovine serum albumin for 30 min and incubated overnight at 4 °C with a rabbit monoclonal antibody against MJFR14-6-4-2. The next day, sections were rinsed in PBS and processed with a rabbit two-step detection kit (PV-9001, ZSGB-BIO, Beijing, China) according to the manufacturer’s instructions. Signal was developed using a 3,3′-diaminobenzidine (DAB) substrate (ZLI-9019, ZSGB-BIO) for 3–5 min under microscopic control. Slides were then washed in tap water, counterstained with hematoxylin, dehydrated through graded ethanol, cleared in xylene, and coverslipped with a permanent mounting medium (cat# E675007-0100, Sangon Biotech, Shanghai, China). Bright-field images of the substantia nigra and surrounding midbrain regions were acquired using a DX150 microscope system (DX150, Shandong Saireedi, Shandong, China) with identical acquisition settings for all groups.

Immunohistochemical detection of p-α-syn was performed on paraffin-embedded brain sections using the same deparaffinization, antigen retrieval, blocking, and DAB detection procedures as described above for MJFR14 immunohistochemistry. For whole-brain mapping, seven representative coronal levels were selected according to their coordinates relative to bregma: +4.28, +2.22, +1.10, −1.82, −2.80, −4.04, and −5.80 mm, based on a previously published study(*47*). At each level, p-α-syn immunoreactivity was evaluated in major PD–related regions, and bright-field images were acquired using the DX150 microscope system with identical acquisition settings for all groups.

#### Quantification of Dopaminergic Neurons and Striatal Terminals

The total number of tyrosine hydroxylase (TH)-positive neurons in the substantia nigra pars compacta (SNpc) and ventral tegmental area (VTA) was estimated using an unbiased stereological approach (*48*). Frozen coronal brain sections (40 μm thick) spanning the rostrocaudal extent of the SNpc (bregma −2.92 mm to −3.52 mm) (*49*) and including the VTA were identified according to The Mouse Brain in Stereotaxic Coordinates, 3rd edition (Paxinos and Franklin, 2007). For each animal, 15 serial sections were collected, and every second section (1-in-2 series; 6 sections in total) was processed for TH immunohistochemistry(*50*).

Sections were then processed for TH staining using the same antigen retrieval, blocking, primary and secondary antibody incubation, and DAB visualization procedures described above. After staining, sections were mounted on gelatin-coated slides, dehydrated through graded ethanol, cleared in xylene, and coverslipped with a permanent mounting medium.

Stereological counting was performed with the optical fractionator probe in Stereo Investigator software (MBF Biosciences, Williston, VT, USA) using a Leica microscope equipped with a motorized stage and a high-resolution digital camera (Leica DFC450, Leica Microsystems). In each section, the anatomical boundaries of the SNpc and VTA were delineated, and TH-positive neurons were sampled using a systematic random sampling design. Stereological parameters, including counting frame size, sampling grid, and guard zone thickness, were kept constant across all animals (*51*). The optical fractionator algorithm was used to estimate the total number of TH-positive neurons in the SNpc and VTA for each animal. All analyses were performed by an investigator blinded to genotype and treatment.

To assess the density of TH-positive fibers in the striatum, bright-field images of TH-stained coronal sections were acquired using a DX150 microscope system. Regions of interest within the striatum were delineated based on anatomical landmarks, and the optical density (OD) of TH immunoreactivity was measured using ImageJ software (NIH, USA). Background signal from an adjacent TH-negative area (corpus callosum) was subtracted (*52*). All images were captured under identical acquisition settings and analyzed using the same thresholding parameters by an investigator blinded to genotype and treatment.

### Nissl, Hematoxylin–Eosin, and Fluoro-Jade C Staining

#### Nissl staining

To assess overall cytoarchitecture and neuronal morphology, Nissl staining was performed on brain sections. Mouse brain sections were first fixed in 70% ethanol and then dehydrated through graded ethanol solutions. Sections were deparaffinized and cleared in xylene, rinsed in distilled water, and incubated in 1% tar violet solution (G1430, Solarbio, Beijing, China) for 30 min at room temperature. After staining, sections were washed with distilled water, briefly differentiated and fixed in 70% ethanol, dehydrated again in graded ethanol, cleared in xylene, and coverslipped with a permanent mounting medium.

#### Hematoxylin–eosin (H&E) staining

To evaluate general tissue morphology and potential systemic pathology, paraffin-embedded sections of brain, colon, heart, liver, spleen, lung, and kidney from WT, S129A, and S129D mice were stained with a hematoxylin–eosin (HE) staining kit (G1120, Solarbio, Beijing, China) according to the manufacturer’s instructions. After staining, sections were dehydrated, cleared, and mounted with a permanent medium. HE-stained sections were examined under a bright-field microscope, and representative images were acquired for qualitative comparison across genotypes.

#### Fluoro-Jade C staining

To detect degenerating neurons, Fluoro-Jade C (FJC) staining was performed on select brain regions using a ready-to-dilute FJC staining kit (BSS-TR-100-FJT, Biosensis, Thebarton, Australia), following the manufacturer’s protocol. Briefly, mounted brain sections were rehydrated, incubated with the FJC staining solution, rinsed, and air-dried before coverslipping with an anti-fade medium (Beyotime, Shanghai, China). FJC-positive cells were visualized with a fluorescence microscope using the FITC-specific filter settings, and images were acquired under identical exposure conditions for all groups. FJC-positive neurons were counted in defined regions of interest using ImageJ and expressed as cell number per mm².

### Immunofluorescence (IF)

#### Double-label immunofluorescence for TH and aggregated α-syn

Double-label immunofluorescence was used to assess the colocalization of aggregated α-syn with dopaminergic neurons. Paraffin-embedded midbrain sections containing the substantia nigra were cut at 4–5 μm and mounted on glass slides. Sections were deparaffinized in xylene, rehydrated through graded ethanol to distilled water, and subjected to heat-induced antigen retrieval in 10 mM sodium citrate buffer (pH 6.0) for 15–20 min. After cooling to room temperature, sections were washed in PBS, permeabilized, and blocked in PBS containing 0.3% Triton X-100 and 3% bovine serum albumin (BSA) for 1 h at room temperature.

Sections were then incubated overnight at 4 °C with a mixture of primary antibodies (table S1) against tyrosine hydroxylase (TH) and aggregated α-syn (5G4 or MJFR14-6-4-2, depending on the experiment). The next day, sections were rinsed in PBS and incubated with appropriate species-specific secondary antibodies conjugated to distinct fluorophores for 1–2 h at room temperature in the dark. Nuclei were counterstained with DAPI. After final washes in PBS, sections were coverslipped with an anti-fade mounting medium. Images were acquired using a Zeiss LSM 880 confocal laser scanning microscope (Carl Zeiss, Oberkochen, Germany) under identical acquisition settings for all groups.

#### Proteinase K treatment and detection of PK-resistant **α**-syn aggregates

To evaluate proteinase K (PK)–resistant α-syn aggregates within dopaminergic neurons, an adjacent paraffin section from each midbrain level was processed in parallel with the double-label immunofluorescence protocol. Sections underwent the same deparaffinization, rehydration, antigen retrieval, and PBS washes as described above. Immediately after antigen retrieval and cooling, one section from each pair was incubated at 37 °C for 30 min in PK buffer containing 10 mM Tris-HCl (pH 7.8), 100 mM NaCl, and 0.1% Nonidet P-40, with proteinase K (EO0491, Thermo Fisher Scientific) diluted to a final concentration of 50 μg/mL, adapted from a previous study(*53*). The paired section was processed without PK and served as a control.

After PK digestion, sections were thoroughly rinsed in PBS to terminate the reaction, then permeabilized and blocked in PBS containing 0.3% Triton X-100 and 3% BSA for 1 h at room temperature. Sections were then incubated overnight at 4 °C with primary antibodies against TH and α-syn. On the following day, sections were washed in PBS, incubated with fluorophore-conjugated secondary antibodies, counterstained with DAPI, and mounted with an anti-fade medium (Beyotime, Shanghai, China), following the same procedure as for the non–PK-treated sections. PK-resistant α-syn immunoreactivity within TH-positive neurons was visualized using the Zeiss LSM 880 microscope. PK-treated and untreated adjacent sections were imaged with identical laser power, detector gain, and pinhole settings and analyzed in parallel to assess the persistence of α-syn signal after PK digestion.

#### TSA-based multiplex immunofluorescence for Lewy body– and Lewy neurite–like pathology

To perform multiplex detection of Lewy body– and Lewy neurite–associated markers in TH-positive neurons of 12-month-old S129A mice, tyramide signal amplification (TSA)–based multiplex immunofluorescence was performed on paraffin-embedded midbrain sections. Sections were deparaffinized, rehydrated, and subjected to antigen retrieval in citrate buffer as described above, followed by permeabilization and blocking in PBS containing 0.3% Triton X-100 and 3% BSA.

Multiplex staining was carried out in sequential rounds. In each round, sections were incubated with a single primary antibody (e.g., TH, aggregated α-syn, and additional Lewy body–associated markers) overnight at 4 °C, followed by incubation with an HRP-conjugated secondary antibody and visualization with a tyramide-based fluorophore working solution according to the manufacturer’s instructions. After each TSA development step, sections were washed thoroughly, and residual HRP activity and bound antibodies were quenched using an appropriate blocking or stripping solution (G1256-50T, Servicebio, Wuhan, China) to prevent cross-reactivity in subsequent rounds. The next primary antibody was then applied, and the staining cycle was repeated until all markers of interest had been labeled.

At the end of the final staining round, sections were counterstained with DAPI, washed, and mounted with an anti-fade medium. Ultrahigh-resolution images of TH-positive neurons containing Lewy body– and Lewy neurite–like inclusions were acquired using a Leica STED FALCON super-resolution fluorescence lifetime imaging system (Stellaris 8 STED, Leica Microsystems, Wetzlar, Germany). STED images were collected with identical excitation, depletion, and detection settings across samples, and z-stacks were acquired for three-dimensional assessment of inclusion morphology and marker colocalization.

The software Imaris (Bitplane, version 10.1.0) was used for three-dimensional reconstruction and visualization of immunofluorescence signals to examine the spatial association of α-syn aggregates with subcellular structures, as described previously(*54*). Briefly, confocal z-stacks (two optical sections spanning 6 μm along the z-axis) containing multichannel signals (DAPI/TH/5G4/Ub/LAMP1/TOM20 or DAPI/TH/MJFR14-6-4-2/TUJ1/Ub/TOM20) were imported into Imaris. After selecting a region of interest (ROI), 3D objects were generated for each channel using the Surfaces or Spots modules, as appropriate, based on the absolute intensity of the corresponding source channel. Thresholds were set to distinguish specific signals from background, and nonspecific objects were manually removed prior to rendering. The resulting 3D reconstructions were used to visualize the subcellular distribution of 5G4- or MJFR14-6-4-2–positive aggregates within TH-positive neurons and their spatial proximity to ubiquitin, LAMP1-positive lysosomal structures, and TOM20-positive mitochondrial structures.

#### HPLC analysis of dopamine and its metabolites

Striatal dopamine and its metabolites were quantified by high-performance liquid chromatography with electrochemical detection (HPLC-ECD). After rapid decapitation, the striatum was dissected on ice, separated from surrounding tissue, and immediately snap-frozen in liquid nitrogen. Before extraction, frozen striatal samples were transferred to pre-chilled microcentrifuge tubes and weighed (JA5003B, Shanghai Yueping Instrument Co., Ltd., Shanghai, China; readability, 0.001 g). Monoamine levels were expressed as micrograms per milligram of wet tissue.

For sample preparation, tissues were homogenized on ice in 150 μL of ice-cold solution A containing 0.4 M perchloric acid. Homogenization was performed by brief ultrasonic disruption on ice (several 1-s pulses with 1-s intervals). Samples were kept on ice and protected from light for 1 h to allow complete protein precipitation, and then centrifuged at 12,000 rpm for 20 min at 4 °C. A defined volume of the supernatant (typically 120 μL) was collected and mixed with half its volume of solution B, a neutralization buffer containing 20 mM potassium citrate, 300 mM dipotassium hydrogen phosphate (K HPO), and 2 mM EDTA in distilled water. After vortexing, the mixture was kept on ice for another 1 h, centrifuged again at 12,000 rpm for 20 min at 4 °C, and the final supernatant (approximately 150 μL) was transferred to HPLC vials and stored at −80 °C in the dark until analysis.

Chromatographic separation was performed on a Waters HPLC system equipped with an autosampler and electrochemical detector (e2695/2475/2465/2998, Waters, Milford, MA, USA) using a reverse-phase C18 analytical column (4.6 × 150 mm). Samples were injected into the column and eluted with a mobile phase consisting of 8% methanol in an aqueous buffer containing 3 mM sodium heptanesulfonate, 100 mM sodium acetate, 85 mM citric acid, and 0.2 mM EDTA, at a constant flow rate according to the manufacturer’s recommendations. Dopamine (DA), 3,4-dihydroxyphenylacetic acid (DOPAC), and homovanillic acid (HVA) were identified by their retention times and quantified by comparison with external standard curves. Final concentrations of DA, DOPAC, and HVA were normalized to original wet tissue weight dissection weights and expressed as μg per mg of wet striatal tissue.

#### Sequential Extraction of Protein Fractions

To evaluate both overall α-syn expression and its distribution between soluble and aggregated pools, brain tissue was processed to obtain (1) total protein lysates, (2) RIPA-soluble fractions, and (3) RIPA-insoluble, urea-solubilized fractions enriched in aggregated proteins (*55*). All procedures were performed on ice or at 4 °C, and all buffers were supplemented with protease and phosphatase inhibitor cocktails (Merck, Darmstadt, Germany).

For total protein extraction, half of one cerebral hemisphere was dissected, weighed, and homogenized in ice-cold RIPA buffer containing 2% SDS. A commercial RIPA lysis buffer (Beyotime, Shanghai, China) was used and supplemented with SDS (Sigma-Aldrich, St. Louis, MO, USA) to a final concentration of 2%. Homogenates were further disrupted by brief sonication and centrifuged at 12,000 rpm for 20 min at 4 °C, and the supernatant was collected as the total protein lysate. These samples were used to assess overall expression levels of α-syn, p-α-syn, and β-Syn by western blotting.

For extraction of RIPA-soluble proteins, the midbrain was rapidly dissected and homogenized in ice-cold standard RIPA buffer by mechanical disruption followed by sonication. Homogenates were centrifuged at 12,000 rpm for 20 min at 4 °C. The resulting supernatant was collected as the RIPA-soluble fraction, representing the detergent-soluble pool of intracellular proteins. The corresponding pellets from this step were retained for subsequent isolation of RIPA-insoluble proteins.

For extraction of RIPA-insoluble (aggregated) proteins, the pellets obtained after RIPA extraction were resuspended in a denaturing buffer containing 8 M urea and 2% SDS (both from Sigma-Aldrich) in Tris-based buffer. Samples were sonicated and vortexed thoroughly to facilitate solubilization of aggregated proteins and then centrifuged again at 12,000 rpm for 20 min at 4 °C. The supernatant was collected as the RIPA-insoluble, urea-solubilized fraction, enriched in aggregated α-syn species, in line with previously described sequential extraction approaches in α-syn research (*56*). Protein concentrations in all fractions were determined by BCA assay, and equal amounts of protein from each fraction were loaded for western blot analysis.

#### Western Blotting

Protein samples from total lysates, RIPA-soluble fractions, and RIPA-insoluble urea–SDS fractions were used for western blot analysis. After sequential extraction, protein concentration in each fraction was determined using a BCA assay according to the manufacturer’s instructions. Equal amounts of protein (typically 15–30 μg per lane) were mixed with SDS sample buffer, heated at 95 °C for 5 min, and separated on 12.5% SDS–polyacrylamide gels (Epizyme Biotech, Shanghai, China). For urea–SDS–solubilized RIPA-insoluble fractions, equal volumes corresponding to comparable tissue equivalents were loaded when accurate protein quantification was not feasible.

Following electrophoresis, proteins were transferred onto polyvinylidene difluoride (PVDF) membranes. Membranes were briefly rinsed in Tris-buffered saline with 0.1% Tween-20 (TBST) and blocked with 5% nonfat milk in TBST for 1 h at room temperature. For the detection of α-syn and p-α-syn, membranes were pre-fixed in 0.4% paraformaldehyde (PFA) in PBS for 30 min at room temperature before the blocking step, and then washed thoroughly in TBST. Membranes were incubated overnight at 4 °C with primary antibodies, followed by incubation with appropriate near-infrared fluorescent secondary antibodies (LI-COR Biosciences, Lincoln, NE, USA) for 1 h at room temperature in the dark.

Fluorescent signals were detected using an Odyssey imaging system (LI-COR Biosciences), and images were acquired with non-saturating exposure settings. Band intensities were quantified using ImageJ software (NIH, USA), with β-actin serving as an internal loading control for normalization. For fractionation experiments, signals from RIPA-soluble and RIPA-insoluble fractions were analyzed in parallel to assess the redistribution of α-syn species between soluble and aggregated pools across genotypes and treatments.

#### Transmission Electron Microscopy (TEM)

Transmission electron microscopy was used to examine ultrastructural changes in dopaminergic neurons within the substantia nigra. Mice were deeply anesthetized and transcardially perfused with ice-cold 0.01 M PBS, followed by a fixative solution containing paraformaldehyde and glutaraldehyde in 0.1 M phosphate buffer. Midbrain blocks containing the substantia nigra were dissected according to stereotaxic coordinates, cut into small pieces (approximately 1 mm³), and post-fixed in the same fixative for several hours at 4 °C.

After rinsing in phosphate buffer, tissue blocks were post-fixed in 1% osmium tetroxide, dehydrated through a graded ethanol series, and infiltrated and embedded in epoxy resin. Semi-thin sections (0.5–1 μm) were first cut and stained with toluidine blue to identify the substantia nigra region and select areas enriched in dopaminergic neurons. Ultrathin sections (approximately 70–90 nm thick) were then cut with an ultramicrotome, collected on copper grids, and contrasted with uranyl acetate and lead citrate.

Ultrathin sections were examined using a TH7700 transmission electron microscope (Hitachi High-Tech, Tokyo, Japan) operated at an appropriate accelerating voltage. Representative images of neuronal somata, axons, and synaptic profiles within the substantia nigra were acquired for qualitative assessment of mitochondrial morphology, intracellular inclusions, and other ultrastructural abnormalities across genotypes and treatment groups.

#### Spatial Transcriptomics

Spatially resolved transcriptomic profiling was performed using the Stereo-seq platform (BGI, Shenzhen, China) to characterize transcriptional alterations in midbrain regions containing dopaminergic neurons. Mice were deeply anesthetized, brains were rapidly removed, embedded in optimal cutting temperature (OCT) compound, and snap-frozen. Coronal cryosections (10 μm thick) at midbrain levels encompassing the substantia nigra and ventral tegmental area were collected onto Stereo-seq chips according to the manufacturer’s instructions, following the standard Stereo-seq protocol described previously(*57*). Sections were briefly warmed to promote tissue adhesion, fixed, permeabilized, and processed for on-chip reverse transcription to capture polyadenylated RNA on the DNA nanoball array. After tissue removal, cDNA was released, amplified, and converted into sequencing libraries. Libraries were sequenced on an MGI DNBSEQ platform with paired-end reads to obtain spatially barcoded transcriptome data for each section.

Raw sequencing data were processed by the service provider using the standard Stereo-seq pipeline to generate gene-by-spot count matrices and corresponding spatial coordinates for each chip. Downstream analysis was performed in Python/R using common single-cell and spatial transcriptomics workflows, with minor adaptations from a previous study(*58*). Briefly, spots or bins with very low transcript counts or detected gene numbers were filtered out, and count data were normalized and log-transformed. Dimensionality reduction and clustering were performed, and spatial domains enriched for dopaminergic neurons and other major cell types were annotated based on canonical marker genes and anatomical location. Spatial expression maps were generated to visualize the distribution of disease-relevant genes and pathways in the midbrain. Differential expression analyses between genotypes and/or age groups were carried out on normalized counts, and significantly altered genes (false discovery rate–corrected p < 0.05) were subjected to gene ontology and pathway enrichment analyses and were used to identify significantly enriched biological processes and pathways, including those related to mitochondrial function, synaptic signaling, and proteostasis.

### Statistical Analysis

All data are presented as mean ± SEM unless otherwise indicated. For animal experiments, n refers to the number of biologically independent mice per group; sample sizes for each assay are provided in the figure legends. Statistical analyses were performed using GraphPad Prism (version 10, GraphPad Software).

For comparisons between two independent groups, two-tailed unpaired t tests were used when data met assumptions of normality and homogeneity of variance. When more than two groups or factors were compared, one-way or two-way ANOVA models were applied as appropriate, followed by Tukey’s multiple comparison tests to correct for post hoc pairwise contrasts. If normality or equal variance assumptions were not satisfied, non-parametric tests (such as the Mann–Whitney U test or Kruskal–Wallis test with Dunn’s post hoc correction) were used instead. Correlation analyses between behavioral, biochemical, and histological readouts were performed using Pearson’s or Spearman’s correlation coefficients, as specified in the figure legends.

For spatial transcriptomic and other high-dimensional datasets, multiple-testing correction was applied using the Benjamini–Hochberg procedure, and genes or pathways were considered significantly altered at a false discovery rate (FDR) < 0.05. For all other experiments, a P value < 0.05 was considered statistically significant. Exact P values, test types, and group sizes are reported in the figure legends. Behavioral testing, image acquisition, and quantitative outcome assessment were performed by investigators blinded to genotype and treatment allocation, with group identities revealed only after completion of outcome assessments.

## Acknowledgments

We thank BioRender (https://biorender.com) for providing the platform to create the schematic diagrams included in this manuscript. JC discloses support by the Senior Research Career Scientist Award (821-RC-NB-30556) from the Department of Veterans Affairs. JC discloses support for the publication of this work from the University of Pittsburgh School of Medicine (the Richard King Mellon Endowed Chair).

## Author contributions

Conceptualization: J.L., J.C.

Methodology: Z.T., Z.X., M.G., H.F., H.Y., Q.S., D.W., F.J., Y.G., H.L., G.L.

Investigation: J.L., J.C., R.L, Z.T., Z.X.

Visualization: J.L., Z.T., Z.X.

Funding acquisition: J.L.

Project administration: J.L.

Supervision: J.L.

Writing – original draft: J.L., Z.T., Z.X.

Writing – review & editing: J.L., J.C., R.L.

## Data and materials availability

The Stereo-seq raw data that supports the findings of this study have been deposited in the Genome Sequence Archive (https://ngdc.cncb.ac.cn/gsa/) under accession number CRA035042 and will be publicly released upon publication.

---

**Table S1.** Antibodies.

| **Name** | **Product ID** | **RRID** | **Host** | **Dilution** |
| --- | --- | --- | --- | --- |
| α-syn | BD Bioscience (610787) | / | Mouse | 1:1000 (WB) |
| α-syn | Cell Signaling Tech<br>(2628) | AB_915792 | Mouse | 1:200 (IF) |
| P-α-syn | Cell Signaling Tech<br>(23706) | AB_2798868 | Rabbit | 1:10000 (WB) |
| P-α-syn | Abcam (ab51253) | AB_869973 | Rabbit | 1:1000 (IHC, IF) |
| β-Syn | Proteintech (10498-1-AP) | AB_2270787 | Rabbit | 1:1000 (WB) |
| TH | Abcam (ab137869) | AB_2801410 | Rabbit | 1:500 (IHC, IF) |
| TH | Millipore (MAB318) | AB_2313764 | Mouse | 1:500 (IF) |
| NeuN | Abcam (ab104224) | AB_10711040 | Mouse | 1:500 (IF) |
| 5G4 | Millipore (MABN389) | AB_2716647 | Mouse | 1:2000 (WB, IF) |
| MJFR14-6-4-2 | Abcam (ab209538) | AB_2714215 | Rabbit | 1:5000 (IHC, IF) |
| LAMP1 | Proteintech (21997-1-AP) | AB_2878966 | Rabbit | 1:1000 (IF) |
| Ub | Proteintech (10201-2-AP) | AB_671515 | Rabbit | 1:500 (IF) |
| TOM20 | Proteintech (11802-1-AP) | AB_2919775 | Rabbit | 1:1000 (IF) |
| TUJ1 | Proteintech (66375-1-Ig) | AB_2814998 | Mouse | 1:500 (IF) |
| β-actin | Huabio (EM21002) | AB_2819164 | Mouse | 1:10000 (WB) |
| β-actin | Huabio (ET1701-80) | AB_2943481 | Rabbit | 1:10000 (WB) |
| Alexa Fluor-488 | Invitrogen (A-11094) | AB_221544 | Rabbit | 1:500 (IF) |
| Alexa Fluor-647 | Invitrogen (A-21235) | AB_2535804 | Mouse | 1:500 (IF) |
| IRDye 680RD | LI-COR Biosciences<br>(926-68070) | AB_10956588 | Mouse | 1:10000 (WB) |
| IRDye 800CW | LI-COR Biosciences<br>(926-32211) | AB_621843 | Rabbit | 1:10000 (WB) |

**Table S2.**
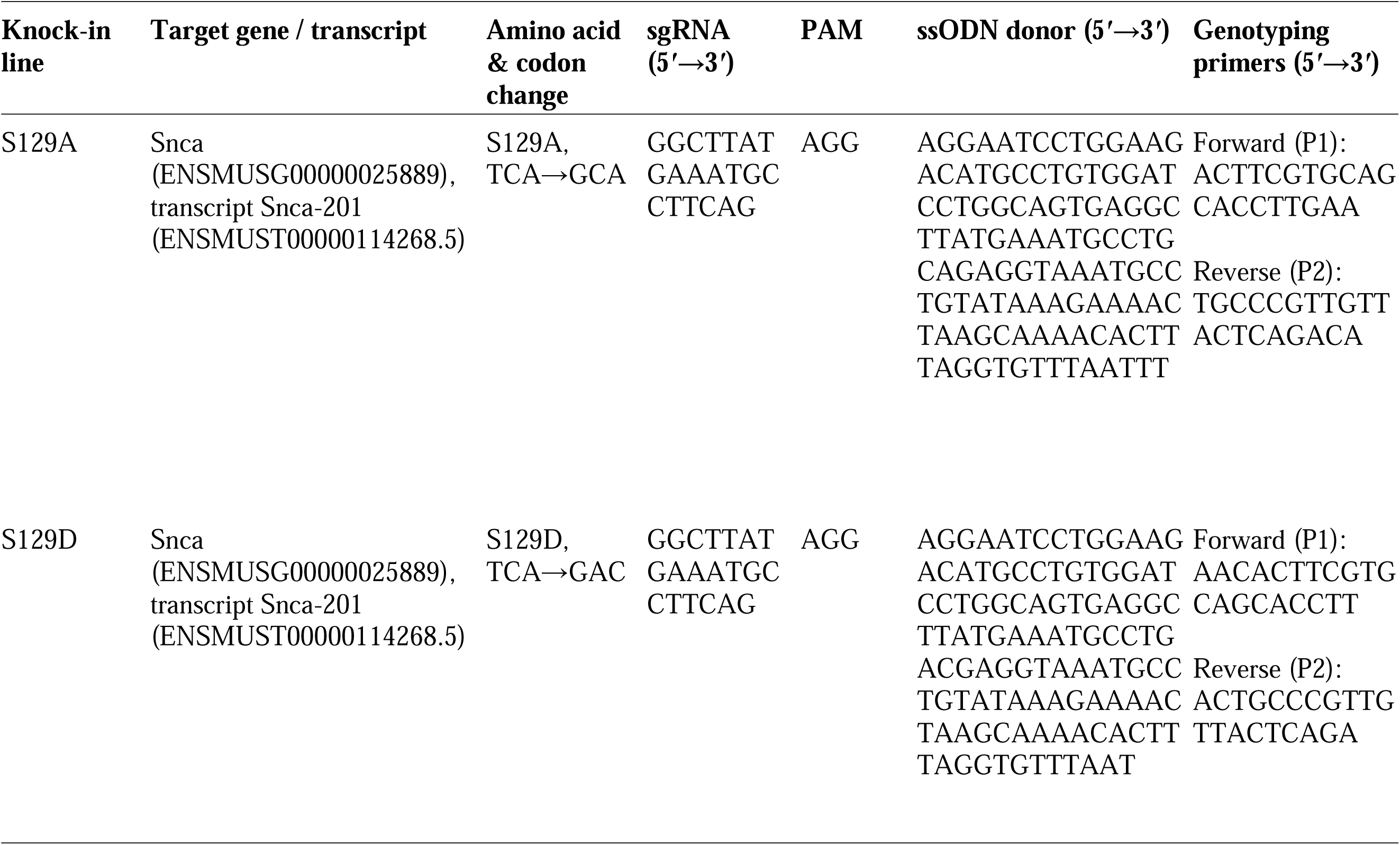
CRISPR/Cas9 reagents used for generation of S129A and S129D knock-in mice.

## Supplemental figure legends

**Fig. S1. Generation and validation of S129A/S129D SNCA knock-in mice, with normal baseline physiological phenotypes.**

(A) Schematic of CRISPR-Cas9-mediated generation of S129A/S129D *SNCA* KI mice: structures of WT, targeted, and point-mutated *Snca* alleles are shown, with homologous recombination and oligonucleotide-directed DNA repair introducing the S129A/D point mutation (exons 5, marked by *). (B) Sequencing validation of S129 site mutations in S129A KI, and S129D KI mice: sequencing traces show the WT (TCA), S129A (GCA), and S129D (GAC) codons. (C–F) Western blot quantification of α-syn, p-α-syn, and β-syn levels (normalized to WT) in brain lysates of WT, S129A KI, S129D KI, α-syn OE and KO mice (n=4–6 mice per group). (G) Representative images of WT, S129A KI, and S129D KI mice at 3 and 12M. (H) Number of weaned pups per litter for WT, S129A KI, and S129D KI breeding pairs (n = 8–10). (I–K) Baseline physiological measurements across 3–12M: food intake percentage (food intake/weight, I), body weight (J), and organ index (ratio of organ weight to body weight) of heart, lung, liver, spleen, and kidney (K) in WT, S129A KI, and S129D KI mice (n = 6–8). (L) H&E staining of major organs (heart, lung, liver, spleen, kidney) in WT, S129A KI, and S129D KI mice, showing normal histological features. All data are presented as mean ± SEM. Quantification analyses (D–F, H) used one-way ANOVA with Tukey’s post hoc test. Analyses of dual-variable groups (I–K) used two-way ANOVA with Tukey’s post hoc test. α-syn, α-synuclein; β-syn, β-synuclein; p-α-syn, phosphorylated α-synuclein; KI, knock-in; OE, overexpression; KO, knockout; WT, wild-type.

**Fig. S2. Consistent loss of TH^+^ and TH^+^NeuN^+^ dopaminergic neurons in SNpc of S129A knock-in mice.**

(A) Immunofluorescence co-staining of SNpc from WT, S129A KI, and S129D KI mice: TH (green) labels dopaminergic neurons, while NeuN (purple) labels all neurons (scale bar: 100 and 20 μm). (B) Quantification of TH^+^ cell counts in SNpc across genotypes (n=6). (C) Quantification of TH^+^NeuN^+^ double-positive cell counts in SNpc across genotypes (n=6). (D) Pearson correlation analysis between TH^+^ cell counts (ratio to WT) and TH^+^NeuN^+^ cell counts (ratio to WT) in SNpc: the positive correlation (r=0.6513, p=0.0034) confirms consistent loss of dopaminergic neurons (both TH^+^ and TH^+^NeuN^+^ counts) in S129A KI mice. All data are presented as mean ± SEM. Quantification analyses (B–C) used one-way ANOVA with Tukey’s post hoc test. Correlation analysis (D) used Pearson correlation. KI, knock-in; NeuN, neuronal nuclei antigen; SNpc, substantia nigra pars compacta; TH, tyrosine hydroxylase; WT, wild-type.

**Fig. S3. Age-dependent dopaminergic system dysfunction in S129A knock-in mice.**

(A–B) TH immunostaining results of the striatum (A, scale bar: 500 μm) and midbrain (B, scale bar: 200 μm) from individual WT, S129A KI, and S129D KI mice across 1, 3, 6, 9, and 12 M; each column displays the staining of a single mouse (labeled with “genotype-age-sample ID”). (C–D) Quantification of TH^+^ cell counts in the left (C) and right (D) VTA of the midbrain across genotypes and ages (n=5). (E) Quantification of TH^+^ fiber density of NAc in the striatum (normalized to WT) across genotypes and ages (n=5). (F–G) Temporal trajectory of TH+ cell counts in SNpc (F) and TH+ fiber density in CPu (G) in S129A KI mice (normalized to WT), showing age-dependent decline. (H–J) Ratios of DA metabolites in brain lysates across genotypes and ages (n=5): DOPAC/DA (H), HVA/DA (I), and (DOPAC+HVA)/DA (J). All data are presented as mean ± SEM. Analyses of dual-variable groups (C–E, H–J) used two-way ANOVA with Tukey’s post hoc test. Analyses of temporal trajectories (F–G) used one-way ANOVA with Tukey’s post hoc test. DA, dopamine; DOPAC, 3,4-dihydroxyphenylacetic acid; HVA, homovanillic acid; KI, knock-in; SNpc, substantia nigra pars compacta; TH, tyrosine hydroxylase; VTA, ventral tegmental area; WT, wild-type.

**Fig. S4. Age-dependent α-syn aggregation and compartmental accumulation in S129A knock-in mice.**

(A) Quantification showing no genotype-specific differences in total α-syn positive area (ratio to WT) in SN sections of WT, S129A KI, and S129D KI mice at 3 and 12 M (n=6). (B) Immunofluorescence staining in 3M α-syn knockout (KO) mice confirms antibody specificity, with no detectable α-syn, 5G4^+^, or MJFR14^+^ signals in SN sections (Scale bars: 20 μm). (C-D) TH/5G4 (C) and TH/MJFR14-6-4-2 (D) co-staining reveals age-dependent accumulation of 5G4^+^ and MJFR14^+^ signals in S129A KI SN neurons at 3, 6, and 12M (scale bars: 100 μm and 10 μm). (E–F) Western blot and quantification demonstrate elevated 5G4^+^ α-syn in the RIPA-insoluble fraction of 12M S129A KI SN, β-actin as loading control (n=6). (G–H) Immunohistochemical staining (G) and quantification (H) confirm increased MJFR14^+^ signal intensity in 3 and 12M S129A KI SN relative to WT (n=6). All data are presented as mean ± SEM. Quantification analyses (A, F, H) used two-way ANOVA with Tukey’s post hoc test. α-syn, α-synuclein; KO, knockout; KI, knock-in; SN, substantia nigra; TH, tyrosine hydroxylase; WT, wild-type.

**Fig. S5. Spatial transcriptomic and single-cell clustering profiling of WT and S129A knock-in mice.**

(A) UMAP plot showing unsupervised clustering of cells (labeled by cluster identity) from merged WT and S129A datasets. (B) UMAP plot of the same cell population, colored by experimental group (WT vs. S129A). (C) Spatial localization of cell clusters in brain spatial transcriptomic sections from WT and S129Amice. (D) Bar plot depicting the proportion of each cell cluster in WT (blue) and S129A (red) samples. (E) Bar plot showing the relative change in cluster proportion (S129A vs. WT); positive values indicate increased proportion in S129A, while negative values indicate decreased proportion. (F) Bar plot displaying the number of upregulated (red) and downregulated (blue) genes per cell cluster in S129A relative to WT. (G) Heatmap of top significant DEGs (log2 fold change) across all cell clusters in S129A vs. WT. All analyses were conducted using standard spatial transcriptomic and single-cell RNA sequencing pipelines. DEGs, differentially expressed genes; UMAP, Uniform Manifold Approximation and Projection; WT, wild-type.

**Fig. S6. Phenotypic and structural analyses of non-motor symptoms in S129A/S129D knock-in mice.** (A) Surface food-seeking latency across genotypes and ages (n=11-19); ns indicates no significant difference. (B) Nissl staining of olfactory bulb layers (GCL, IPL, MCL, EPL, GL; scale bar: 100 μm) in WT, S129A KI, and S129D KI mice. (C) Quantification of Nissl^+^ cell density in each olfactory bulb layer (normalized to WT; n=6). (D) Fecal wet weight across genotypes and ages (n=5-6). (E) Colon outer longitudinal muscle layer thickness across genotypes (n=3). (F) Colon immune infiltration score across genotypes (n=3). (G) Immunofluorescence staining of colon tight junction proteins (ZO-1: red; Claudin-5: green; DAPI: blue; scale bar: 20 μm) across genotypes. (H-I) Quantification of ZO-1 (H) and Claudin-5 (I) fluorescence intensity in colon tissue (n=3). (J–L) Open field test metrics: total movement distance (J), movement time (K), and mean speed (L) across genotypes and ages (n=11-16). (M) Nissl staining of hippocampal subregions (CA1, CA2, CA3, DG; scale bar: 200 and 50 μm) across genotypes. (N) FJC staining of hippocampal subregions (scale bar: 100 μm) across genotypes. All data are presented as mean ± SEM. Analyses for A, C, D, J-L used two-way ANOVA with Tukey’s post hoc test; analyses for Fig. S6E, F, H-I used one-way ANOVA with Tukey’s post hoc test. EPL, external plexiform layer; FJC, Fluoro-Jade C; GCL, granule cell layer; GL, glomerular layer; IPL, internal plexiform layer; KI, knock-in; MCL, mitral cell layer; WT, wild-type.

**Fig. S7. p-α-syn antibody specificity validation and SNpc p-α-syn expression in WT, S129A, and S129D mice.** (A) p-α-syn immunostaining of brain sections from α-syn KO mice (scale bars: 500 and 100 μm); no p-α-syn signal was detected, confirming antibody specificity. (B) Immunofluorescence co-staining (DAPI: blue; TH: pink; p-α-syn: yellow) of SNpc in 3M WT mice (scale bars: 100 and 10 μm); p-α-syn co-localized with TH-positive dopaminergic neurons. (C) Immunofluorescence co-staining (DAPI: blue; TH: pink; p-α-syn: yellow) of SNpc in 3M S129A KI and S129D KI mice (scale bars: 100 μm and 10 μm); no p-α-syn signal was observed in either genotype. KI, knock-in; KO, knockout; SNpc, substantia nigra pars compacta; TH, tyrosine hydroxylase; WT, wild-type.

**Fig. S8. S129A/S129D homozygous and heterozygous knock-in mice exhibit no significant motor or cognitive behavioral abnormalities at 3 months.** (A) Schematic of the experimental design: the study included mice of five genotypes—WT, S129A homozygotes (S129A^+^/^+^), S129A heterozygotes (S129A^+^/^−^), S129D homozygotes (S129D^+^/^+^) and S129D heterozygotes (S129D^+^/^−^)—with behavioral assessments (covering olfactory, gastrointestinal, motor, and cognitive functions) scheduled at 3, 6, and 9 M of age. (B) Relative change rate of surface food-finding test across genotypes at 3 M (n=10-17). (C–D) Relative change rate of pole test (C) and rotarod test (D) across genotypes at 3 M (n=10-18). (E) Relative change rate of Y-maze test across genotypes at 3 M (n=10-17). All data are presented as mean ± SEM. ns indicates no significant difference. Analyses used one-way ANOVA with Tukey’s post hoc test. KI, knock-in; WT, wild-type.

**Fig. S9. S129A KI mice with or without nigral AAV-mediated S129D intervention show no significant motor or cognitive impairments at 3 months.** (A) Schematic of the experimental design: WT and S129A homozygous (S129A^+^/^+^) mice received targeted injection of AAV-S129D or empty AAV-Vector into the SN at 1 M of age. Behavioral assessments (covering olfactory, gastrointestinal, motor and cognitive function) were scheduled at 3, 6, and 9 M of age. (B) Finding time of surface food-finding test across groups at 3 M. (C) Descent time of pole test. (D) Latency to fall of rotarod test across groups at 3 M. (E) Spontaneous alternation rate of Y-maze test across groups at 3 M. All data are presented as mean ± SEM. ns indicates no significant difference. Analyses used one-way ANOVA with Tukey’s post hoc test. AAV, adeno-associated virus; SN, substantia nigra; WT, wild-type.

**Fig. S10. S129A KI mice treated with Minzasolmin show no significant motor or cognitive impairments at 3 months.** (A) Schematic of the experimental design: WT and S129A homozygous (S129A^+^/^+^) mice were treated with Minzasolmin (an α-syn aggregation inhibitor) or Capisol (vehicle) at 1 M of age. Behavioral assessments (covering olfactory, gastrointestinal, motor, and cognitive functions) were scheduled at 3, 6, and 9 M of age. (B) Finding time of surface food-finding test across groups at 3 M. (C) Descent time of pole test across groups at 3 M. (D) Latency to fall of rotarod test across groups at 3 M. (E) Spontaneous alternation rate of Y-maze test across groups at 3 M. All data are presented as mean ± SEM. ns indicates no significant difference. Analyses used one-way ANOVA with Tukey’s post hoc test. Minzasolmin, α-syn aggregation inhibitor; Capisol, vehicle; KI, knock-in; WT, wild-type.

**Fig. S11. Graphical Abstract: Disease progression and pathogenesis in S129A α-syn knock-in mice.** Disease progression in S129A α-syn KI mice occurs in three temporal stages over 3, 6, 9 and 12 M: pre-motor stage (3 M) with olfactory and gastrointestinal dysfunction, motor-dominant stage (6 M) with motor deficits, and advanced stage (9-12 M) with cognitive impairment. Within SNpc DA neurons, unphosphorylated S129A α-syn forms aggregates that induce downstream pathological dysfunctions (mitochondrial, ubiquitination, and synapse dysfunction), ultimately leading to progressive neurodegeneration and SNpc DA neuron loss. DA, dopaminergic; KI, knock-in; SNpc, substantia nigra pars compacta.

