## Supplementary figures and images for "Ser129 Phosphorylation Paradox: Non-phosphorylated α-Synuclein Drives Parkinson’s Disease-like Pathogenesis *in Vivo*"

### Fig. S1

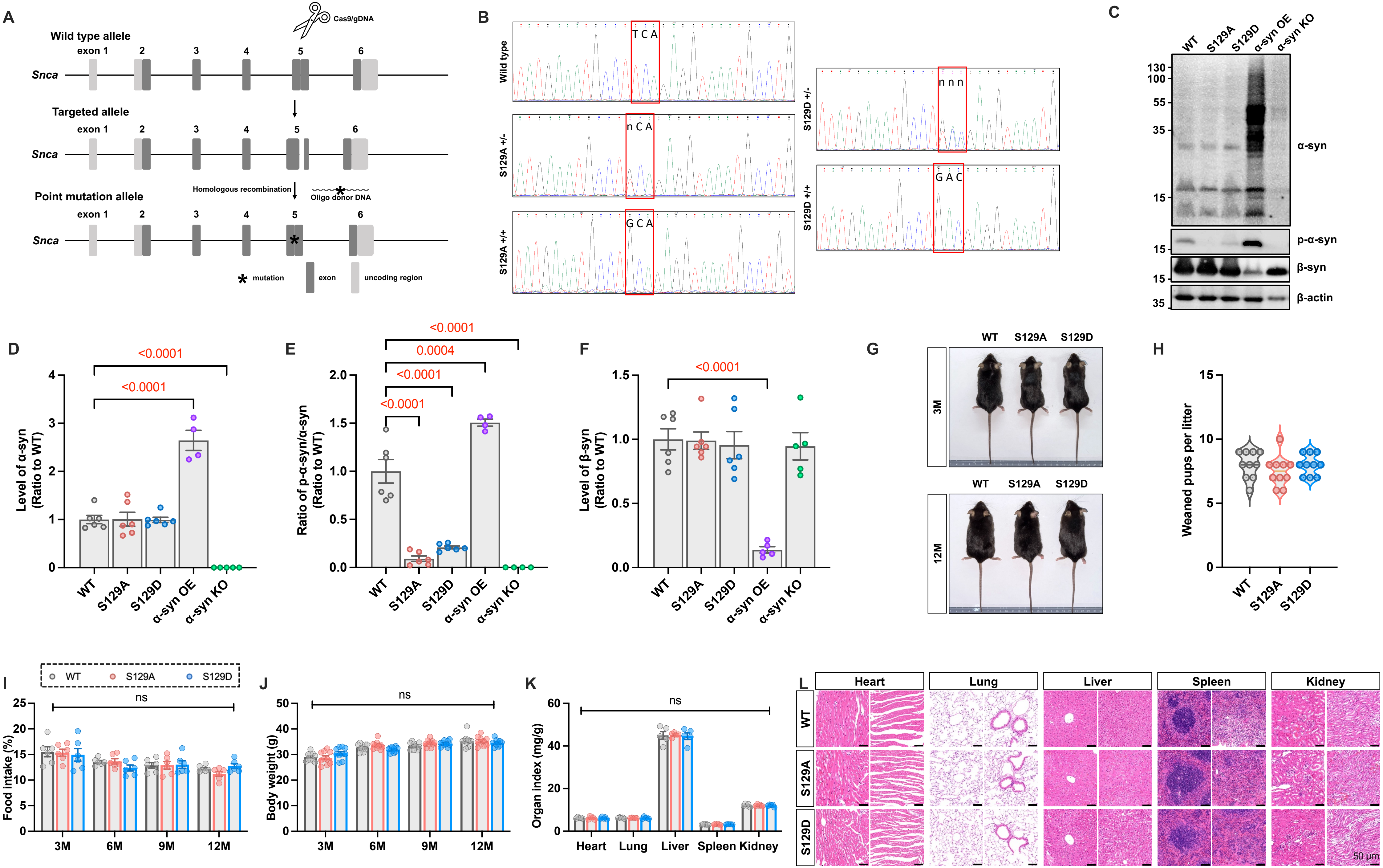

### Fig. S2

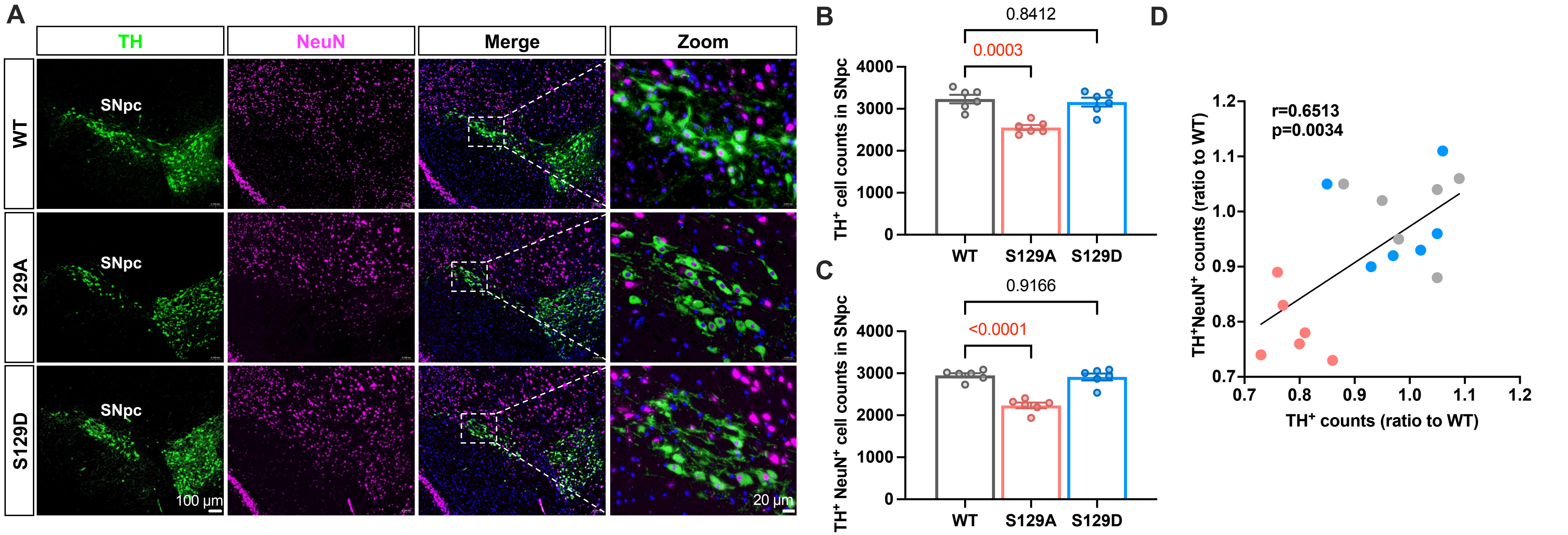

### Fig. S3

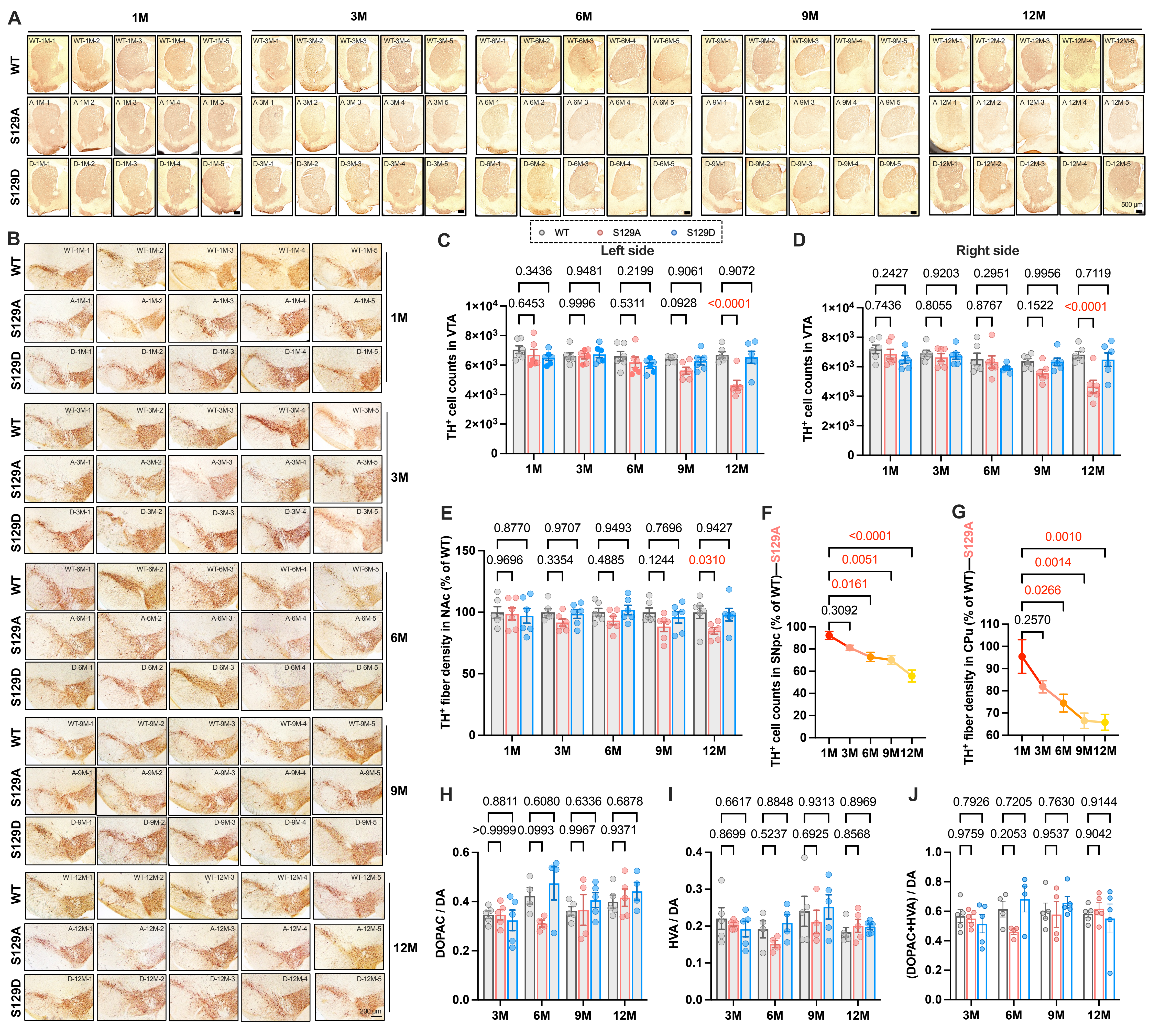

### Fig. S4

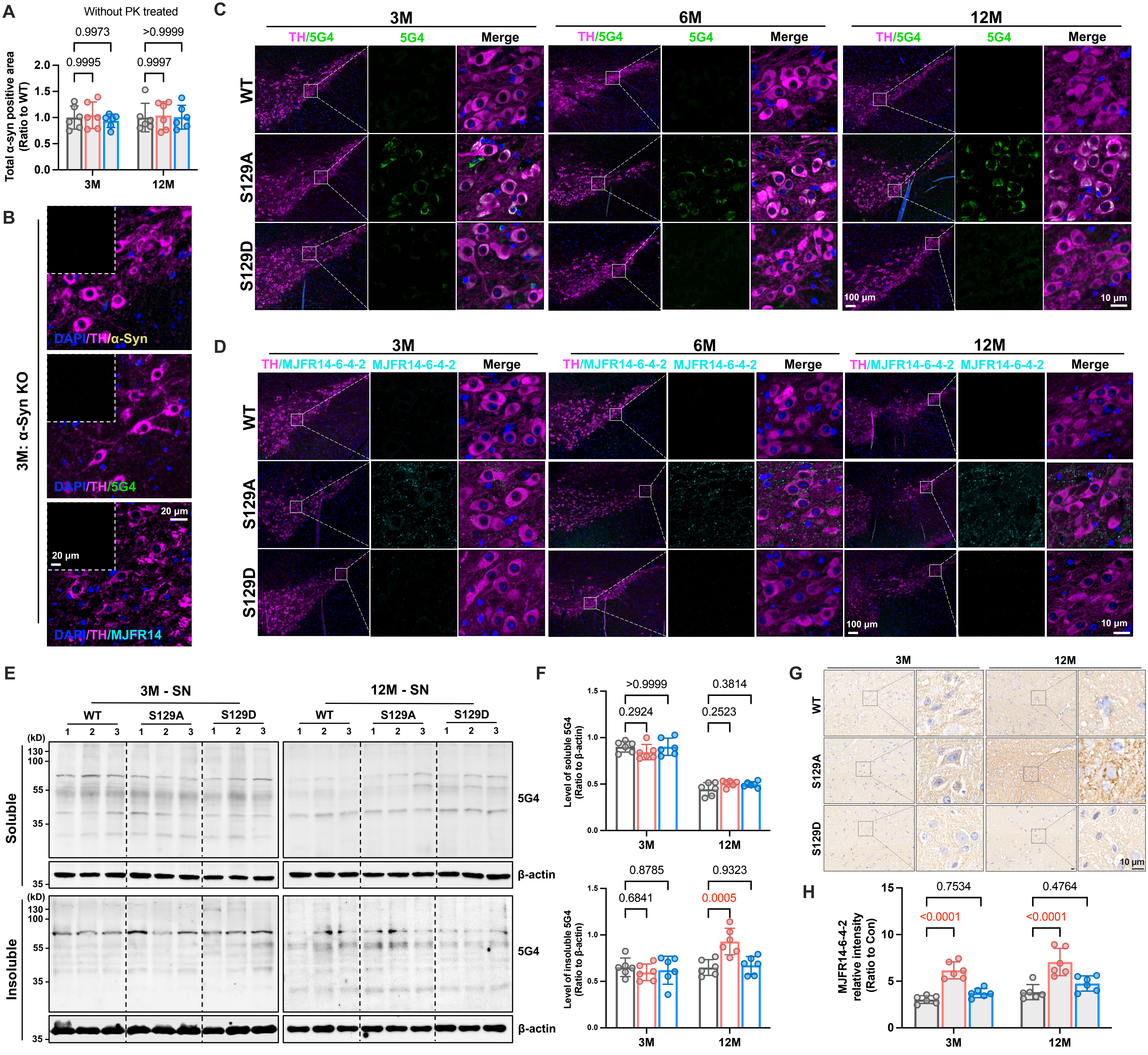

### Fig. S5

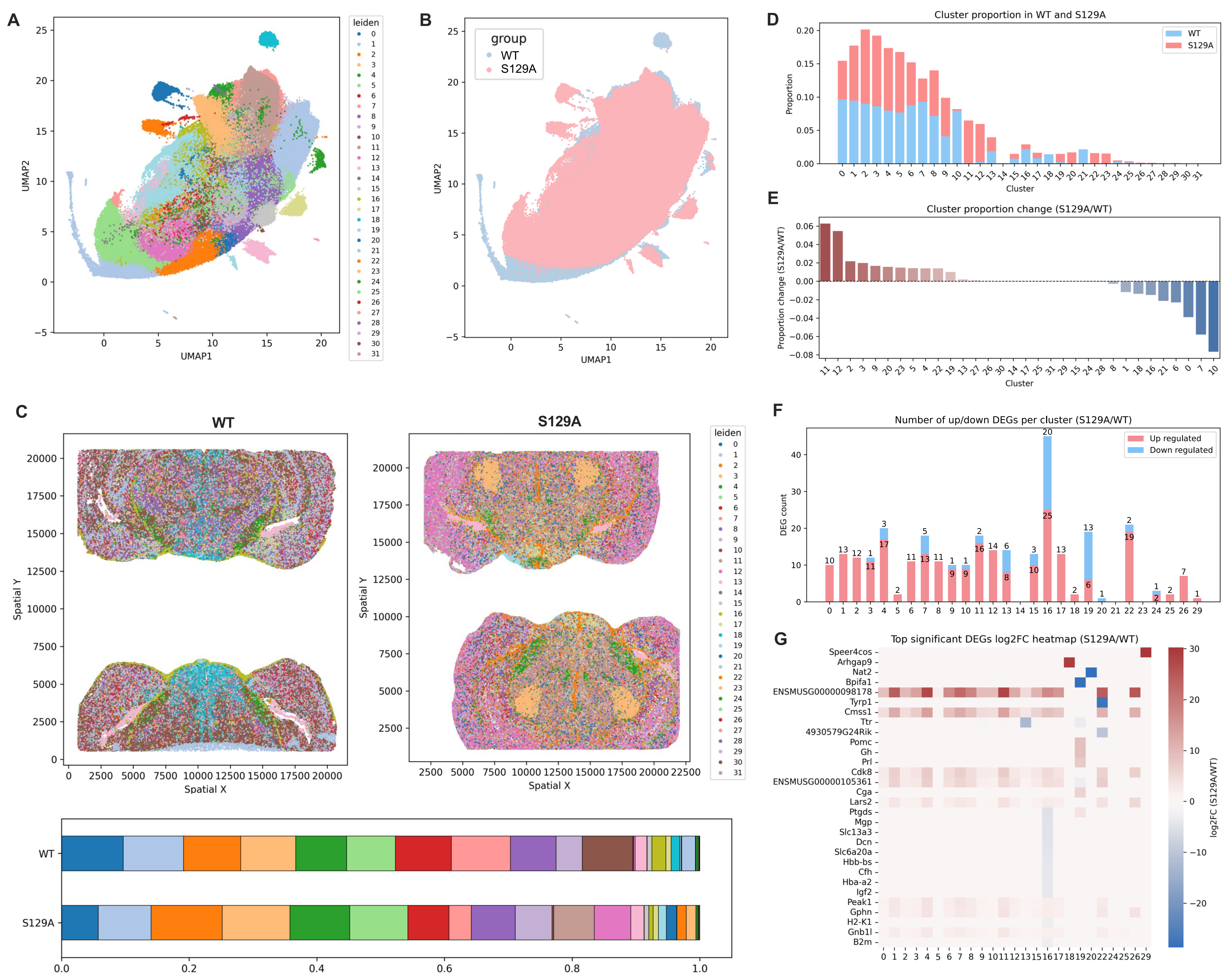

### Fig. S6

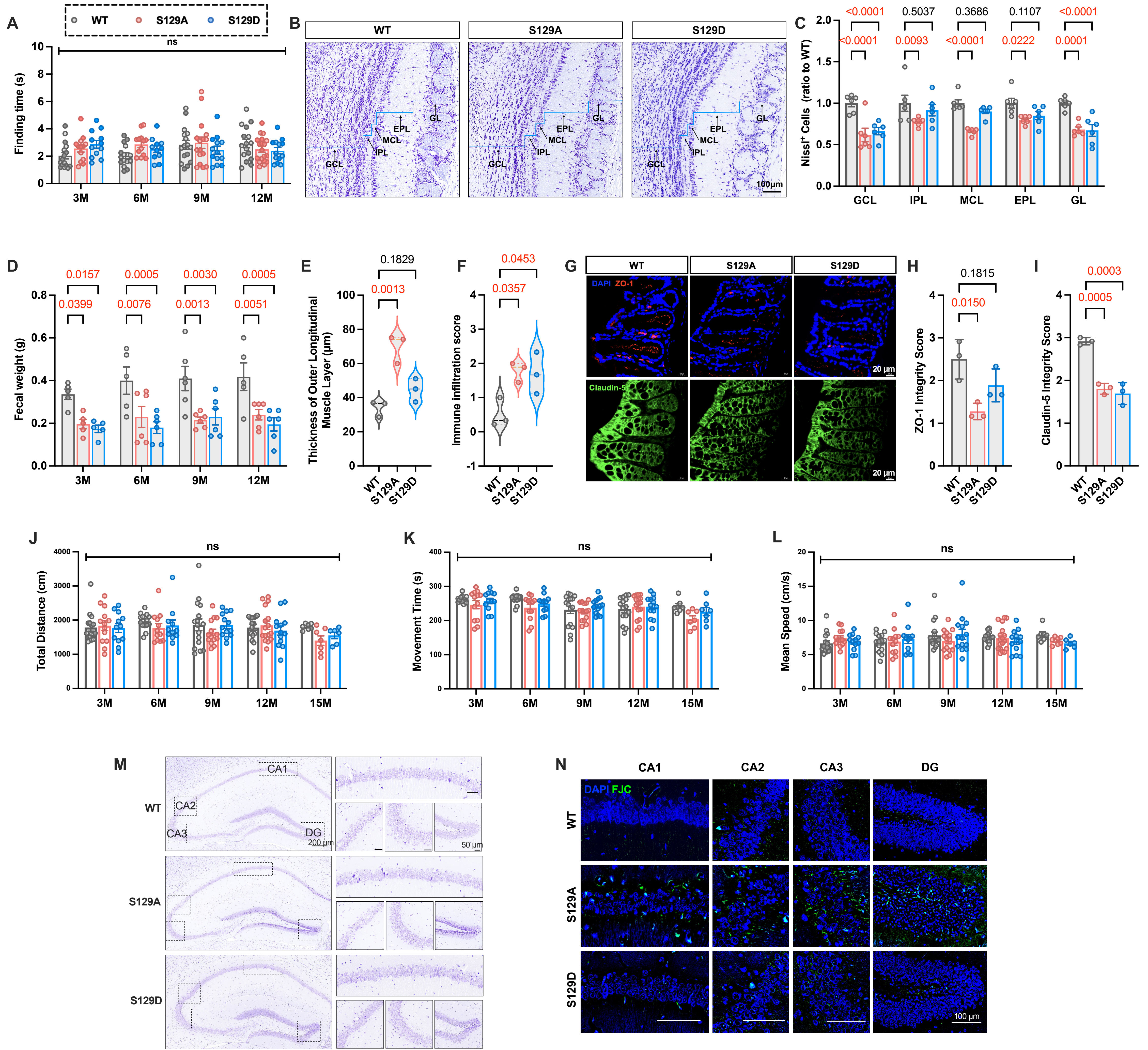

### Fig. S7

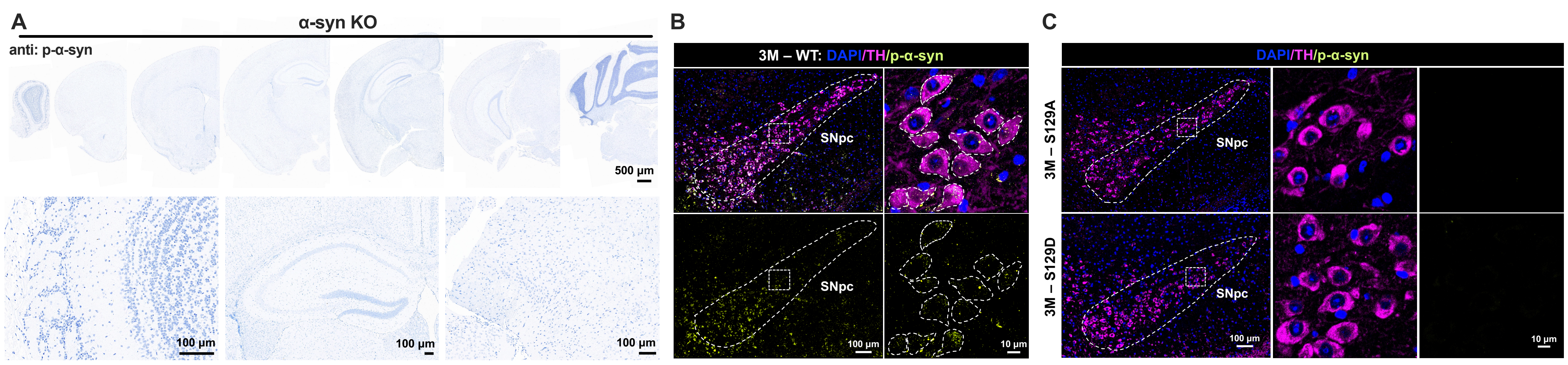

### Fig. S8

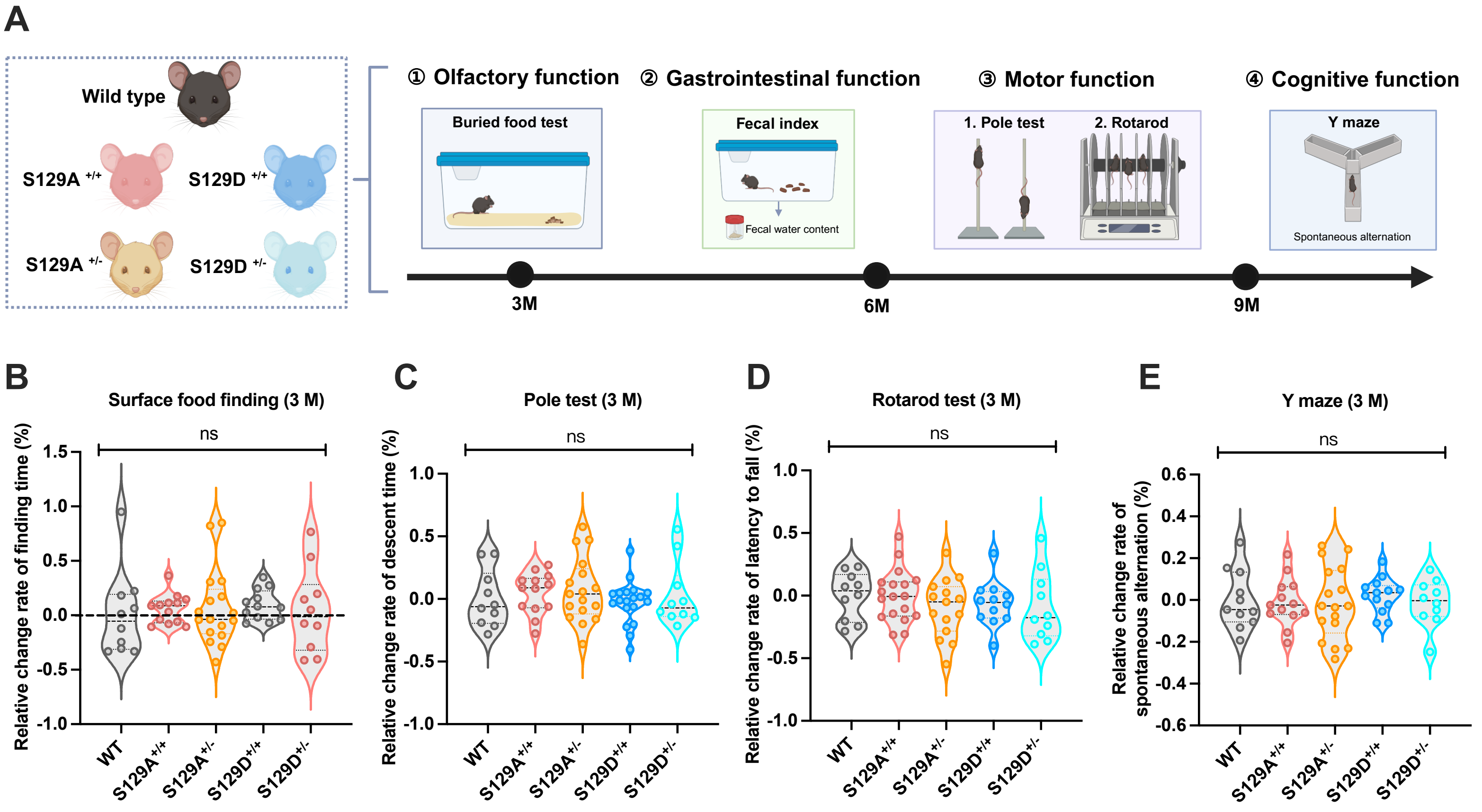

### Fig. S9

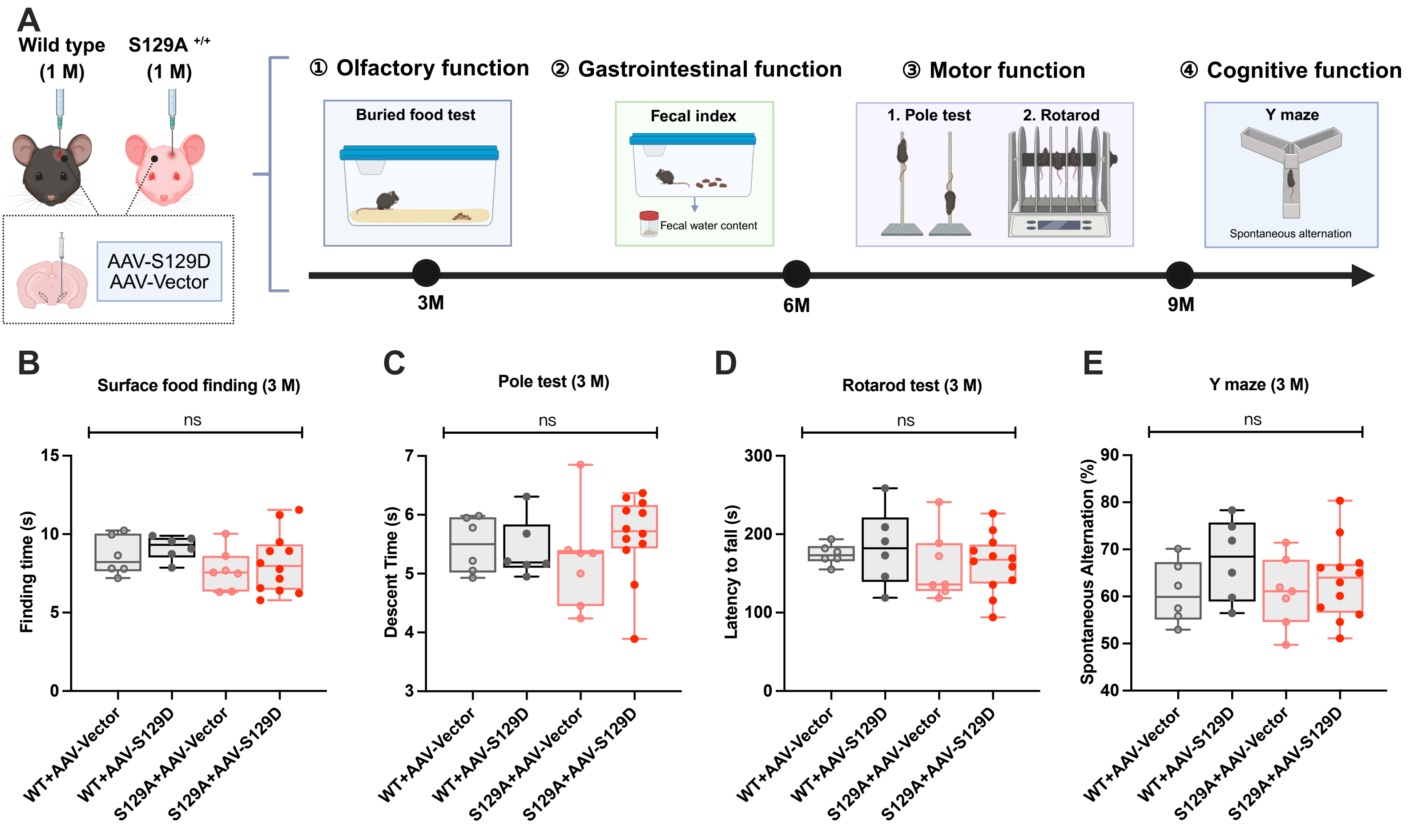

### Fig. S10

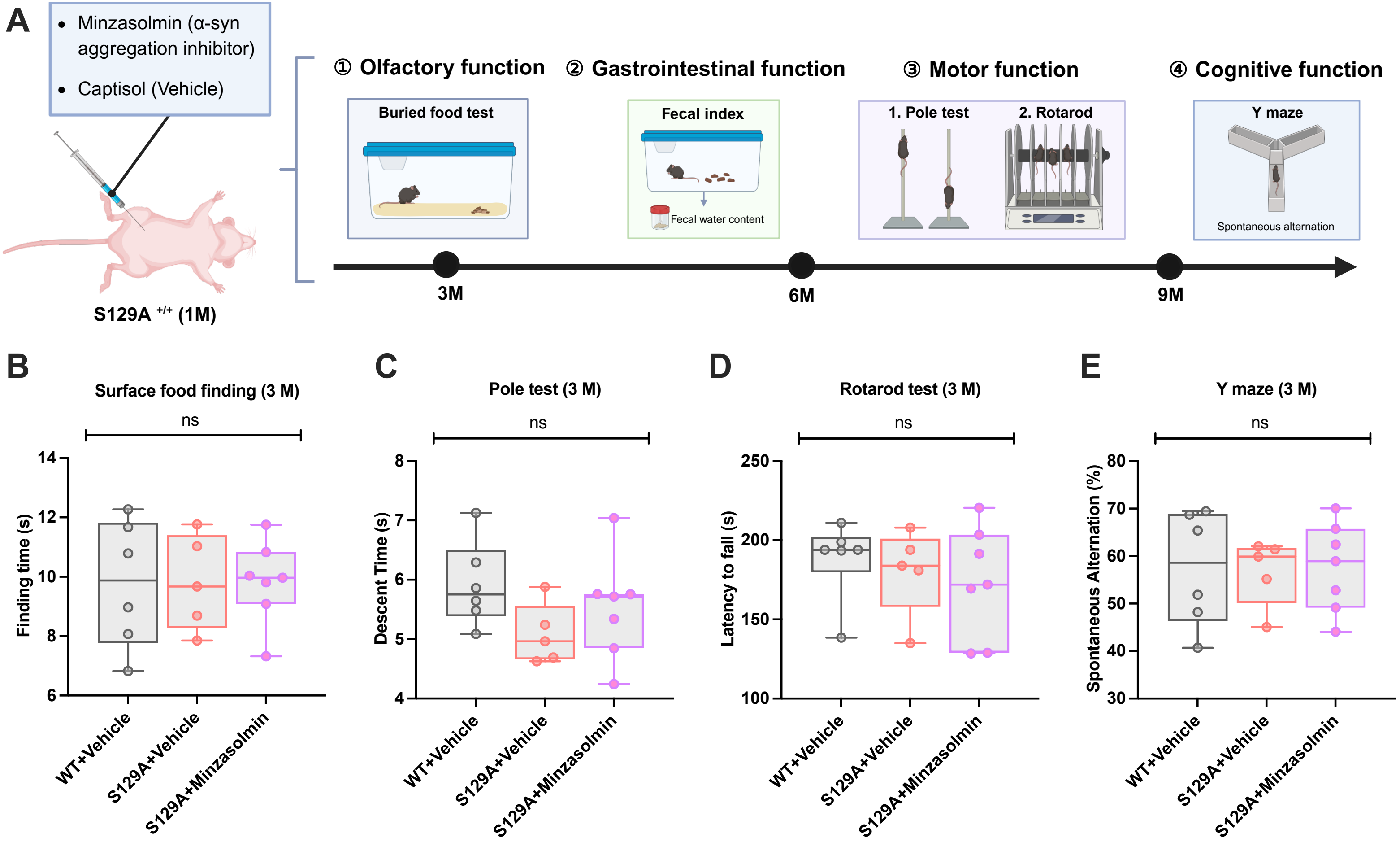

### Fig. S11

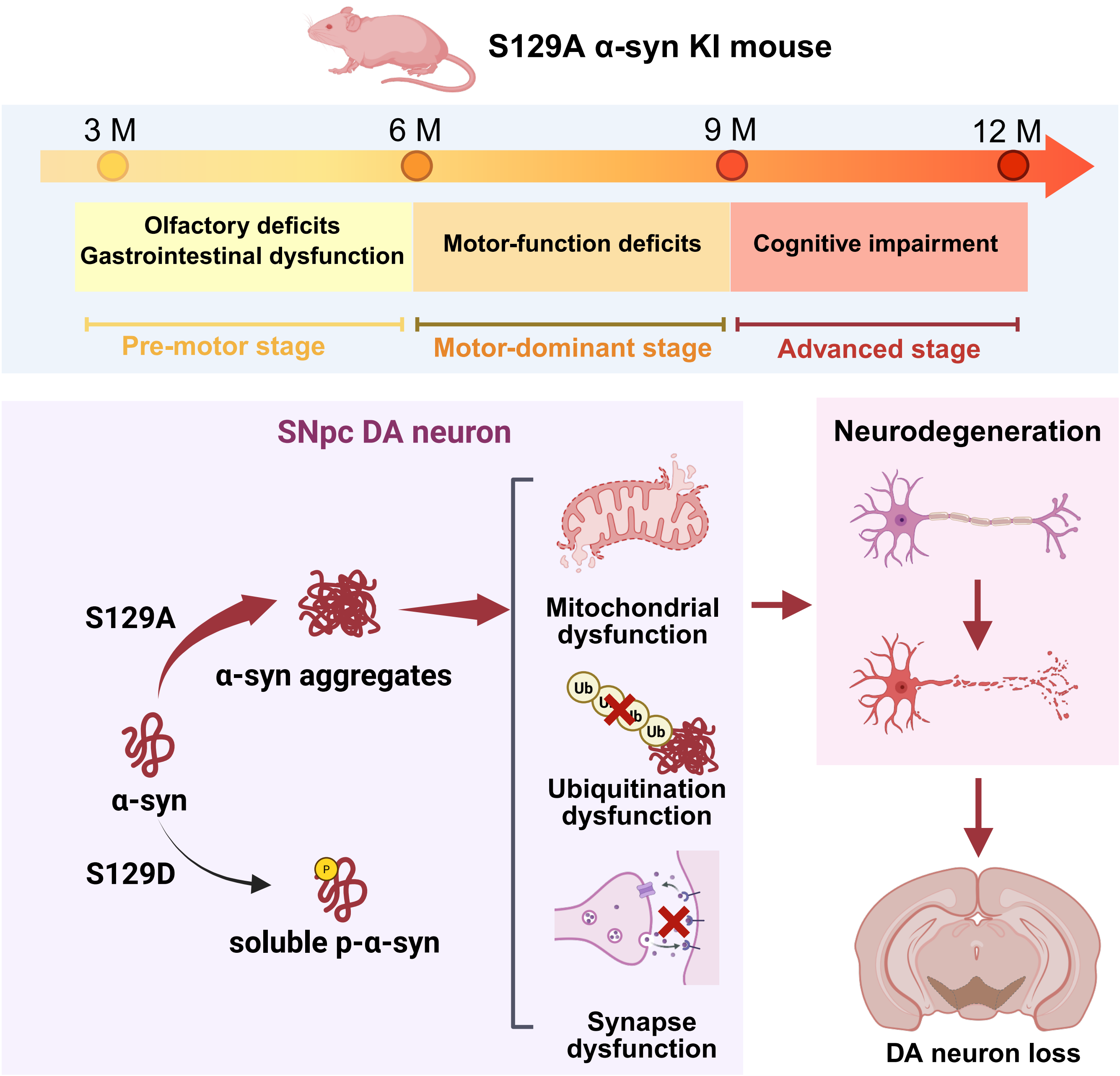
